# A developmental switch to facilitation shapes synaptic plasticity at neuron-OPC synapses in white matter

**DOI:** 10.64898/2026.03.31.715637

**Authors:** Bartosz Kula, Ting-Jiun Chen, Balint Nagy, Anahit Hovhannisyan, David Terman, Wenjing Sun, Maria Kukley

## Abstract

Neuronal circuits rely on precisely timed synaptic transmission and plasticity, which are established through activity-dependent maturation during development. While these processes are well characterized at neuronal synapses, far less is known about how synaptic communication between neurons and glial cells develops. Pyramidal cortical neurons project axons through white matter where they release glutamate ectopically along their shafts and form glutamatergic synapses with oligodendrocyte precursor cells (OPCs). The functional maturation of these neuron-glia connections remains unknown.

Here, using single-cell electrophysiology combined with computational modelling, we show that neuron–OPC synapses in the mouse corpus callosum undergo a pronounced developmental transformation in short-term synaptic plasticity. During the first two postnatal months, these synapses switch from strong synaptic depression to facilitation. This transition is accompanied by a shortening of synaptic delay and a reduction in asynchronous glutamate release, indicating an increase in temporal precision of neurotransmitter signalling in white matter. Computational modelling suggests that both pre- and postsynaptic changes may underlie this functional maturation.

Taken together, our findings demonstrate that neuron-OPC synapses in white matter are not static but undergo developmental transition towards facilitation and temporally precise transmission that parallels the maturation trajectory of classical neuronal synapses in cortical grey matter. These results identify neuron-glia synapses in white matter as dynamic elements of developing neural circuits, and suggest that synaptic release machineries at axonal shafts in white matter and synaptic boutons in grey matter mature in a similar fashion.

## 1. Introduction

Oligodendrocyte precursor cells (OPCs), also known as NG2 cells, are widely distributed in the grey and the white matter of the central nervous system (CNS). Originally considered simply as progenitors of oligodendrocytes (OLs), OPCs came to light upon the discovery that they receive direct glutamatergic and GABAergic synaptic input from neurons in various CNS regions in animals and humans, including the white matter (Kula et al., 2019). Similar to conventional neuronal synapses, neuron-OPC synapses involve the presynaptic vesicular release machinery and the postsynaptic neurotransmitter receptors, and neurotransmission at these synapses occurs on a rapid (milliseconds) time-scale. Neuron-OPC synaptic signaling is thought to be involved in activity-dependent myelination, and may modulate OPC proliferation, migration, and differentiation (Chen et al., 2018; Nagy et al., 2017).

A key phenomenon at neuronal synapses is synaptic plasticity: when neurons fire repetitively, prior synaptic activity modulates subsequent synaptic efficacy, on a short (short-term plasticity, STP) or long (long-term-plasticity, LTP) timescale. STP, rapid and transient change in synaptic efficacy, is thought to be related to the moment-to-moment processing of neuronal information that may be useful in perception and decision-making (Arguello and Gogos, 2012). STP undergoes significant changes with brain maturation, a process that represents a crucial aspect of brain development. If maturation-dependent dynamics of synaptic plasticity is disturbed, this may result in impaired complex information processing within neuronal networks, leading to behavioral and cognitive alterations. For example, reduced STP has been suggested to contribute to the mechanisms of schizophrenia, autism spectrum disorders, Fragile X syndrome, and Alzheimer’s disease (Arguello and Gogos, 2012; Crabtree and Gogos, 2014; Deng et al., 2011; Feng et al., 2003; Giovedi et al., 2014; Lee et al., 2012).

Cortical glutamatergic synapses in the mouse mature during the first four to five weeks after birth (Etherington and Williams, 2011), and during this time-period STP switches from strong depression to weaker depression or even facilitation (Feldmeyer and Radnikow, 2009). Glutamatergic pyramidal neurons located in cortical layers II/III, layer V and layer VI project through the corpus callosum where they release glutamate along their shafts and form glutamatergic synapses with OPCs. Intriguingly, these neuron-OPC synapses, deep in the white matter, also undergo STP (Nagy et al., 2017). But the information about the origin of STP at neuron-OPC synapses (presynaptic vs postsynaptic) is very sparse, and the dynamic maturation-dependent changes of STP are not known. In the present study, we combined experimental and computational approaches to address these questions.

In our experimental approach, we performed patch-clamp recordings from callosal OPCs in brain slices and stimulated callosal axons repetitively in three age-groups of mice: 8-10 days of age (P10 group), 19-22 days of age (P20 group), and 50-53 days of age (P50 group). We selected these age groups for two major reasons: (1) they correspond to distinct postnatal stages of callosal axons development: establishment of innervation between contra-lateral synaptic targets (P10 group), fine-tuning of callosal projections (P20 group), and fully established callosal projections (P50 group) (De Leon Reyes et al., 2020; Mizuno et al., 2010); and (2) they correspond to the distinct postnatal stages of callosal myelination: no/little myelination (P10 group), peak of myelination (P20 group), and slow steady-state level of myelination (P50 group). We analyzed presynaptic and postsynaptic properties of neuron-OPC synapses and STP across these three age-groups.

In our computational approach, we established passive OPC models for each of the three age groups using reconstructed OPC morphology data (Kula and Kukley, 2025), and simulated the STP we observed from experimental recordings using either presynaptic or postsynaptic mechanisms.

Together, our experimental and computational approaches provide new insights into the developmental dynamics of STP at neuron-OPC synapses in the corpus callosum. This knowledge is essential for understanding how OPCs integrate into neuron-glia networks, respond to activity-dependent signals, and process synaptic information for their own development and function. Given that synaptic communication is a potential route through which neuronal activity regulates OPC behavior, aberrant plasticity at neuron-OPC synapses may contribute to OPC dysfunction and impaired myelination during neurological and psychiatric disorders such as multiple sclerosis, schizophrenia, and major depressive disorder (Fields, 2008; Haroutunian et al., 2014; Mitew et al., 2014). Understanding the origin and the mechanisms of this plasticity may reveal novel therapeutic targets to promote re-myelination or restore normal network function.

## 2. Results

### 2.1. Passive membrane properties and expression of K^+^ and Na^+^ channels in callosal OPCs change during animal maturation

OPCs usually show a characteristic electrophysiological profile of voltage-gated currents which distinguish them from neurons and other types of glia in brain slices, and the passive membrane properties of OPCs change with age (Kukley et al., 2007; Larson et al., 2016). We verified the age-dependent dynamics of those parameters in our three groups of animals because they are of key importance for computational modeling.

First, we investigated passive membrane properties of OPCs including the resting membrane potential (V_rest_), the membrane resistance (R_m_), and the cell capacitance (C_m_). We found that those parameters changed drastically during postnatal development. The R_m_ was the highest in OPCs of P10 group (2245 ± 185.80 MΩ), lower in P20 group (588.40 ± 45.78 MΩ), and the lowest in P50 group (179.50 ± 11.71 MΩ), with all of the groups differing statistically significantly: P10 vs P20, p=1.50E-11; P10 vs P50, p=4.90E-14; P20 vs P50, p=8.82E-13 (Figure 1A). The V_rest_ was more hyperpolarized in OPCs from older animals compared to younger ones: -77.88 ± 2.818 at P10, -86.55 ± 0.978 mV at P20, -91.80 ± 1.097 mV at P50 (P10 vs P20, p=6.81E-03; P10 vs P50, p=2.01E-04; P20 vs P50, p=1.57E-03) (Figure 1B). Increased expression of leak K^+^-channels in OPCs of more mature animals underlies changes in V_rest_ and R_m_ (Pivonkova et al., 2024). Higher K^+^ leak conductance shifts V_rest_ toward the K^+^ equilibrium potential and lowers the R_m_ by increasing baseline membrane permeability. Similar mechanisms are well established in neurons, where the background K^+^ currents, often mediated by two-pore domain (K2P/KCNK) channels, constitute a major determinant of V_rest_ and neuronal excitability (Enyedi and Czirjak, 2010; Goldstein et al., 2001). These channels generate voltage-independent leak conductance that stabilizes the negative V_rest_ and oppose depolarization. The C_m_ was lower in OPCs of younger animals compared to older mice: 18.97 ± 0.70 pF at P10, 27.18 ± 0.93 at P20, and 26.76 ± 1.49 pF at P50 (P10 vs P20, p=3.10E-10; P10 vs P50, p=3.76E-05; P20 vs P50, p=0.82) (Figure 1C). Increase in the C_m_ indicates that OPCs in older animals have larger surface area, in line with our recent observations showing that the total length of all OPC processes increases during callosal development, with an increase being detectable already during the third postnatal week (Kula and Kukley, 2025).

**Figure 1:**
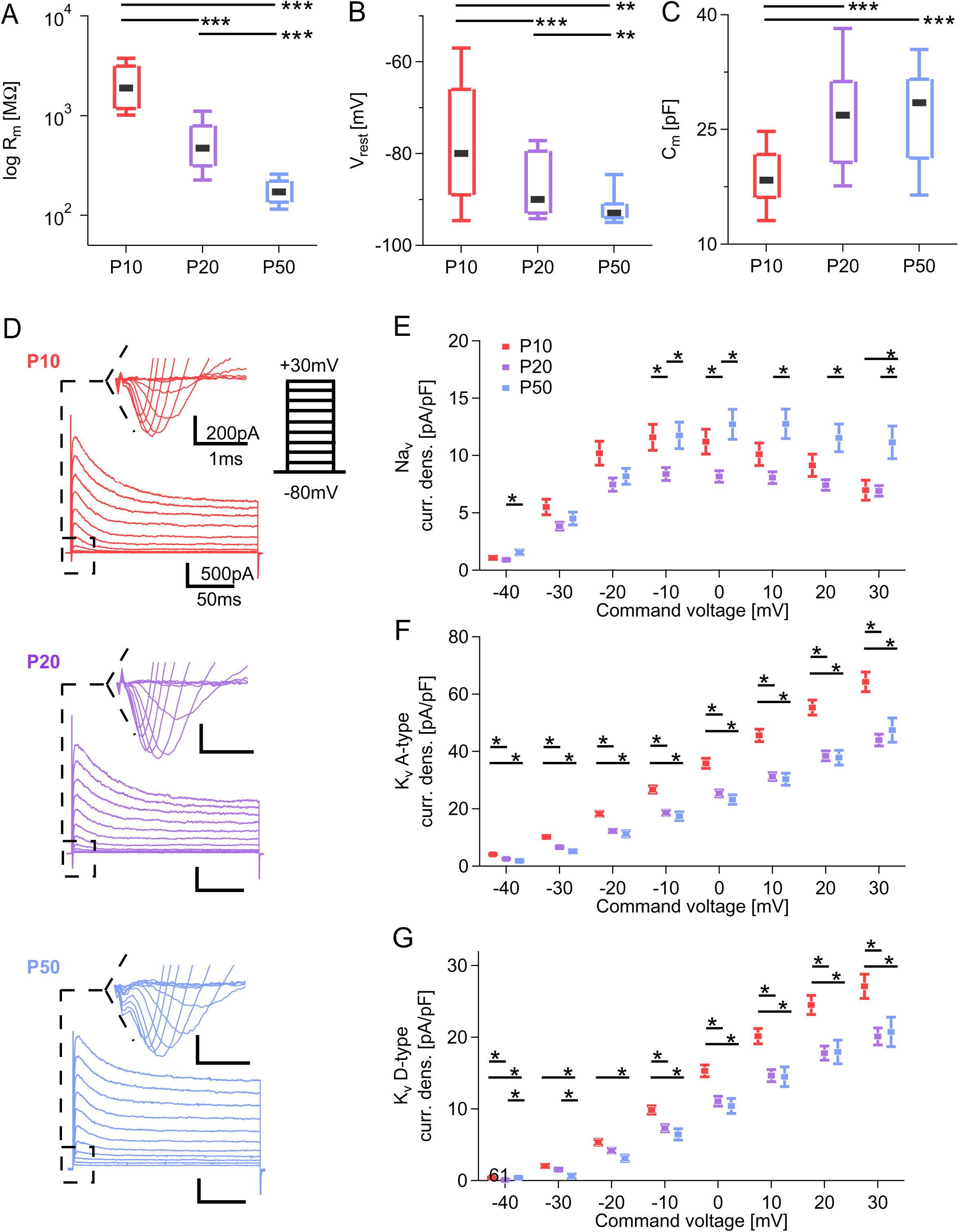
Passive membrane properties and expression of K^+^ and Na^+^ channels in callosal OPCs change during animal maturation. **(A)** : Membrane resistance (R_m_) of OPCs in animals of three age-groups. Box plots represent 25^th^ to 75^th^ percentiles, whiskers represent 10^th^ and 90^th^ percentiles and the black bar is the median. Black horizontal bars link significantly different groups. * = p<0.05; ** = p<0.01; *** = p<0.001. P10 group: n = 46 cells from 38 animals; P20: n = 72 cells from 70 animals; P50: n = 23 cells from 23 animals. **(B)** : Resting membrane potential (V_rest_) of OPCs in animals of three age-groups. P10 group: n = 25 cells from 21 animals; P20: n = 59 cells from 53 animals; P50: n = 20 cells from 20 animals. Box plots as in (A). **(C)** : Membrane capacitance (C_m_) of OPC in animals of three age-groups. P10 group: n = 47 cells from 38 animals; P20: n = 83 cells from 74 animals; P50: n = 35 cells from 32 animals. Box plots as in (A). **(D)** : Representative examples of OPC current responses to 11 incremental (increment = 10mV), 200 ms long command voltage steps, from the holding potential V_h_ = -80 mV. Red: P10 group; purple: P20 group; blue: P50 group. The examples represent recorded raw traces without leak subtraction. The insets highlight fast activating Na^+^ current, characteristic for OPCs. In the inset, the leak current was subtracted. **(E)** : Density of Na^+^-current in OPCs from three age-groups of animals. Each marker represents the group mean ± SEM. The leak current was subtracted from all traces, and for each OPC, currents were normalized on its capacitance. Black horizontal bars link significantly different groups. * = p<0.05; ** = p<0.01; *** = p<0.001. P10 group: n = 45 cells in 36 animals; P20: n = 77 cells from 70 animals; P50: n = 28 cells from 28 animals. **(F)** : Density of A-type K^+^-current in OPCs from three age-groups of animals. Each marker represents the group mean ± SEM. The leak current was subtracted from all traces, and for each OPC, currents were normalized on its capacitance. Black horizontal bars link significantly different groups. * = p<0.05; ** = p<0.01; *** = p<0.001. P10 group: n = 46 cells in 36 animals; P20: n = 84 cells from 70 animals; P50: n = 30 cells from 30 animals. **(G)** : Density of steady-state K^+^-current in OPCs from three age-groups of animals. Each marker represents the group mean ± SEM. The leak current was subtracted from all traces, and for each OPC, currents were normalized on its capacitance. Black horizontal bars link significantly different groups. * = p<0.05; ** = p<0.01; *** = p<0.001. P10 group: n = 46 cells in 36 animals; P20: n = 84 cells from 70 animals; P50: n = 29 cells from 30 animals.

To study the functional voltage-gated K^+^ and Na^+^ channels in OPCs, we performed whole-cell voltage-clamp recordings, applied series of depolarizing voltage steps (11 steps with +10mV increments) from the holding potential (V_h_) of -80 mV (Figure 1D), and analyzed the amplitude of the K^+^ and Na^+^ currents (Figure 1E-G). Leak currents were subtracted before the analysis (see Materials and Methods).

The density of voltage-gated Na^+^-currents in OPCs showed differences between the age-groups at more depolarized potentials, with minor differences between the P10 and P20 groups (at V_step_ to -10mV and 0mV; p=0.037 to 0.026), largest differences between the P20 and P50 groups (consistently at V_step_ potentials between -10mV and +30mV; p=0.0053 to 0.037), and very minor differences between P10 and P50 groups (only at V_step_ to +30mV; p=0.034), (Figure 1E).

The density of voltage-gated A-type K^+^-currents was considerably and consistently higher in P10 OPCs compared to P20 (p=1.42E-06 to 2.93E-04) and to P50 animals (p=3.84E-06 to 6.42E-03), at all tested potentials from -40mV to +30 mV, while the P20 and P50 groups did not differ significantly (p=0.064 to p=0.88), (Figure 1F).

The differences between the age-groups in the density of D-type K^+^-currents in OPCs showed a pattern similar to those of A-type K^+^-currents. OPCs in P10 mice had the highest density of D-type K^+^-currents and tested as significantly different from OPCs in P50 throughout the whole range of tested voltage steps (V_h_=-40mV to +30mV; P10 vs P50: p=3.00E-04 to p=0.020), (Figure 1G). The P10 and P20 groups were consistently different in voltage ranges of V_h_=-10mV to V_h_=+30mV (Figure1G; P10 vs P50: p=2.47E-04 to 0.042) However, as with A-type K^+^-currents, differences between P20 and P50 were minor and observed only at V_step_ between -40 mV and -30 mV (p=0.020, p=0.048, respectively), (Figure 1G).

Taken together, these results show that A-type and D-type K^+^-channel density undergoes marked age-dependent down-regulation in callosal OPCs during postnatal development. The A-type and D-type voltage-gated K^+^ currents are also well described in neurons, where they regulate excitability. A-type currents, mediated primarily by K_v_4 family channels, delay spike initiation and shape dendritic integration, whereas D-type currents, often associated with K_v_1 channels, contribute to spike repolarization and firing adaptation (Falk et al., 2003; Jerng et al., 2004; Mitterdorfer and Bean, 2002; Storm, 1988). In many neuronal populations, the density of transient K^+^ channels increase during maturation, contributing to reduced membrane resistance and stabilization of neuronal firing properties.

### 2.2. STP at neuron-OPC synapses in the corpus callosum changes during postnatal development and may be heterogeneous between the OPCs

STP is a rapid reversible change in synaptic strength occurring over milliseconds to seconds during ongoing activity. To study STP at neuron-OPC synapses in the corpus callosum, we recorded AMPAR-mediated excitatory synaptic currents (EPSCs) in callosal OPCs in response to electrical stimulation of callosal axons with trains of 20 pulses at two different frequencies, 25 Hz and 100 Hz. We have chosen those frequencies because they are widely used for probing STP at neuronal synapses.

Synaptic response of OPCs to repetitive axonal stimulation consisted of two components: (1) phasic EPSC, a response which occurred immediately following an action potential in the axons, and (2) a barrage of small-amplitude EPSCs which were not time-locked to the action potential and continuing for tens of milliseconds after the phasic response (Figure 2A). The two components of synaptic release have also been observed at many neuronal synapses (Kaeser and Regehr, 2014; Kavalali, 2015).

**Figure 2:**
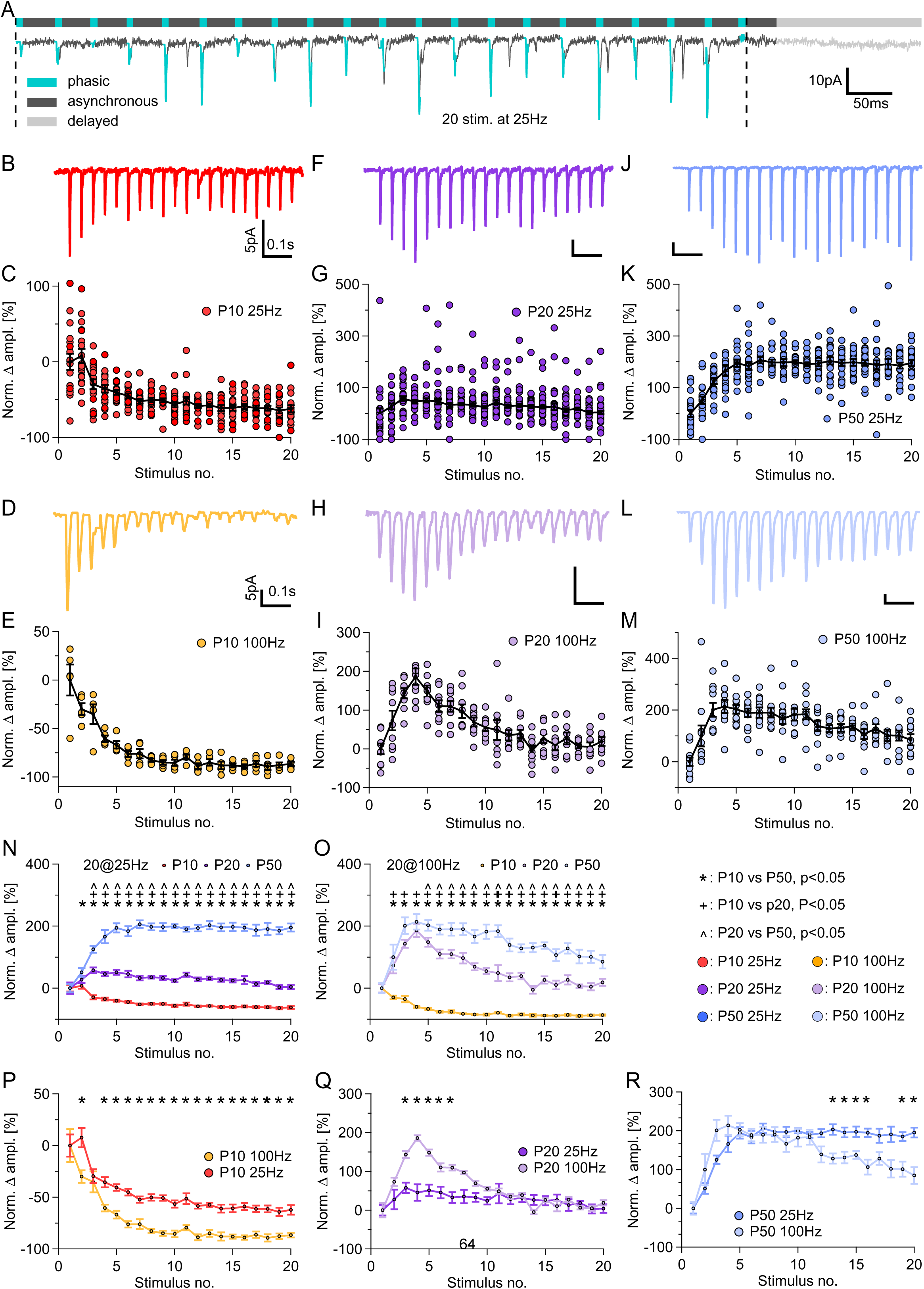
Developmental switch from short-term depression to short-term potentiation at neuron-OPC synapses in the corpus callosum. **_(A)_** : Top: a bar indicating time-periods when phasic (turquoise; 5ms after stimulation onset), asynchronous (black; 35ms after stimulation onset), and delayed (grey; after the cessation of stimulation) release of glutamate occur at neuron-OPC synapses during stimulation of callosal axons with 20 pulses at 25 Hz. Bottom: a representative trace recorded from a callosal OPC (V_h_ = -80 mV) in response to the train stimulation of the callosal axons, P50 group. Dashed lines mark the beginning and end of the stimulation period. **(B)** : Representative example of a current response of the callosal OPCs to the train stimulation of the callosal axons with 20 pulses at 25 Hz in P10 group. The shown trace is an average of 20-60 recorded traces. V_h_ = -80 mV. **(C) :** Dynamics of the phasic EPSCs amplitude during the train. Each circle represents a normalized average of the EPSCs recorded in a single OPC over 20-100 train stimulation trials. Black line with circles represents the group mean ± SEM; n = 15 cells from 14 animals. **(D-E)**: As **(B-C)**, but for the train stimulation of the callosal axons with 20 pulses at 100 Hz; n = 14 cells from 10 animals. **(F-G)**: As **(B-C)**, but for the P20 group; n = 23 cells from 18 animals. **(H-I)**: As **(F-G)**, but for the train stimulation of the callosal axons with 20 pulses at 100 Hz; n = 19 cells from 14 animals. **(J-K)**: As **(B-C)**, but for P50 group; n = 19 cells in 16 animals. **(L-M)**: As **(J-K)**, but for the train stimulation of the callosal axons with 20 pulses at 100 Hz; n = 9 cells from 9 animals. **(N)** : Summary of **(C)**, **(G)**, and **(K)**: STP in OPCs triggered by the stimulation with 20 stimuli at 25 Hz in three age-groups. Each circle represents the group mean ± SEM. P10 in red, P20 in purple, P50 in blue. * = p<0.05, P10 vs P50; ^+^ = p<0.05, P10 vs P20; ^#^ = p<0.05, P20 vs P50. **(O)** : Summary of **(E)**, **(I)**, and **(M)**: STP in OPCs triggered by the stimulation with 20 stimuli at 100 Hz in three age-groups. Each circle represents the group mean ± SEM. P10 in orange, P20 in lime, P50 in purple. * = p<0.05, P10 vs P50; ^+^ = p<0.05, P10 vs P20; ^#^ = p<0.05, P20 vs P50. **(P)** : Summary of **(C)** and **(E)**: STP in OPCs triggered by the stimulation with 20 pulses at 25 or 100 Hz in the P10 group. * = p<0.05. **(Q)** : Summary of **(G)** and **(I)**: STP in OPCs triggered by the stimulation with 20 pulses at 25 or 100 Hz in the P20 group. * = p<0.05. **(R)** : Summary of **(K)** and **(M)**: STP in OPCs triggered by the stimulation with 20 pulses at 25 or 100 Hz in the P50 group. * = p<0.05.

First, we focused on the amplitude of the phasic EPSCs. We considered an EPSC as phasic if its onset occurred no later than 5 ms after a stimulus (unless otherwise indicated). All currents with an onset beyond 5 ms were considered non-phasic and were analyzed later (see below). We found that the STP differed between the age-groups and was dependent on the stimulation frequency. We also noted heterogeneity of STP between the OPCs.

#### STP at neuron-OPC synapses changes during postnatal development

In P10 animals, we observed short-term synaptic depression during the trains: upon 25 Hz stimulation, the phasic EPSC amplitude briefly potentiated (+7.71 ± 9.38 %) and decreased after the second stimulus in the train. By the last stimulus, EPSC amplitude decreased by 62.23 ± 42.66% (Figure 2B-C). The decline was more prominent during the 100 Hz stimulation, where no potentiation was observed, and the amplitudes decreased sharply after the first stimulus, reaching an 86.62 ± 2.20% reduction by the end of the train (Figure 2D-E).

In P20 animals, brief initial potentiation occurred during the first 3-4 stimuli of the train: by 57.61 ± 9.29% and 186.09 ± 21.17%, at peak, at frequencies 25 and 100 Hz, respectively (Figure 2F-I). Subsequently, the phasic EPSC amplitude was declining, although by the end of train the amplitudes still remained slightly potentiated and were higher than the initial amplitude by +3.97 ± 6.62% at 25 Hz and by +18.84 ± 11.31% at 100 Hz (Figure 2F-I).

In P50 animals, upon 25 Hz stimulation, we observed strong synaptic potentiation with continuous increase in the amplitude reaching a plateau around the 7^th^ stimulus. At the plateau, the amplitude was 206.18 ± 12.38% higher than the initial value (Figure 2J-K). Upon 100 Hz stimulation, STP showed a biphasic time-course: the phasic EPSC amplitude steeply increased to +213.91 ± 24.93% of the initial value at the peak but, after a short plateau, started to decrease although remained strongly potentiated (by +84.98 ± 21.88 %) till the end of the train (Figure 2L-M).

Our statistical analysis showed that for the 25 Hz stimulation, the EPSC amplitudes in OPCs of P10 animals were significantly different from those in P20 (p=1.91E-12 to p=6.87E-07) and P50 animals (p<1.00E-15 to p=0.042) throughout the entire train, while significant difference in the phasic EPSC amplitudes between P20 and P50 animals was observed only from the third stimulus in the train (p<1.00E-15 to p=3.11E-05) (Figure 2N). For the 100 Hz stimulation, the EPSC amplitudes in OPCs of P10 animals were also significantly different from P20 (p=9.22E-09 to p=3.73E-04) and P50 animals (p=3.73E-04 to p=0.037) throughout the entire stimulation train, but the significant difference between P20 and P50 groups was observed only from the tenth stimulus (p=3.70E-04 to p=0.044) (Figure 2O). In addition to comparing the differences between the groups, we also compared the differences between the two stimulation paradigms (i.e. 20 pulses at 25 Hz vs 20 pulses at 100 Hz) within the groups. We found that the response patterns tested as having different progression: p=2.64E-04 to 0.018 at P10, p=0.0014 to p=0.034 at P20, and p=5.54E-06 to p=0.044 at P50 (Figure 2P-R).

#### OPCs exhibit heterogeneity in STP profiles

Several recent studies indicated that OPCs represent a heterogeneous population of cells (Pivonkova et al., 2024; Spitzer et al., 2019). We were wondering whether this heterogeneity is also reflected in the synaptic STP profiles. To address this question, we divided OPCs into seven categories, based on the shape and the best possible fit to the STP profile (linear, polynomial, or exponential) during the 25 Hz train stimulation (see Methods). We defined two extreme categories in which the amplitude of the phasic EPSCs either depressed (category 1) or potentiated (category 7) very quickly forming a plateau during the time-course of the train (Figure 3A), and five other categories representing response patterns transitioning between those two extremes (Figure 3A).

**Figure 3:**
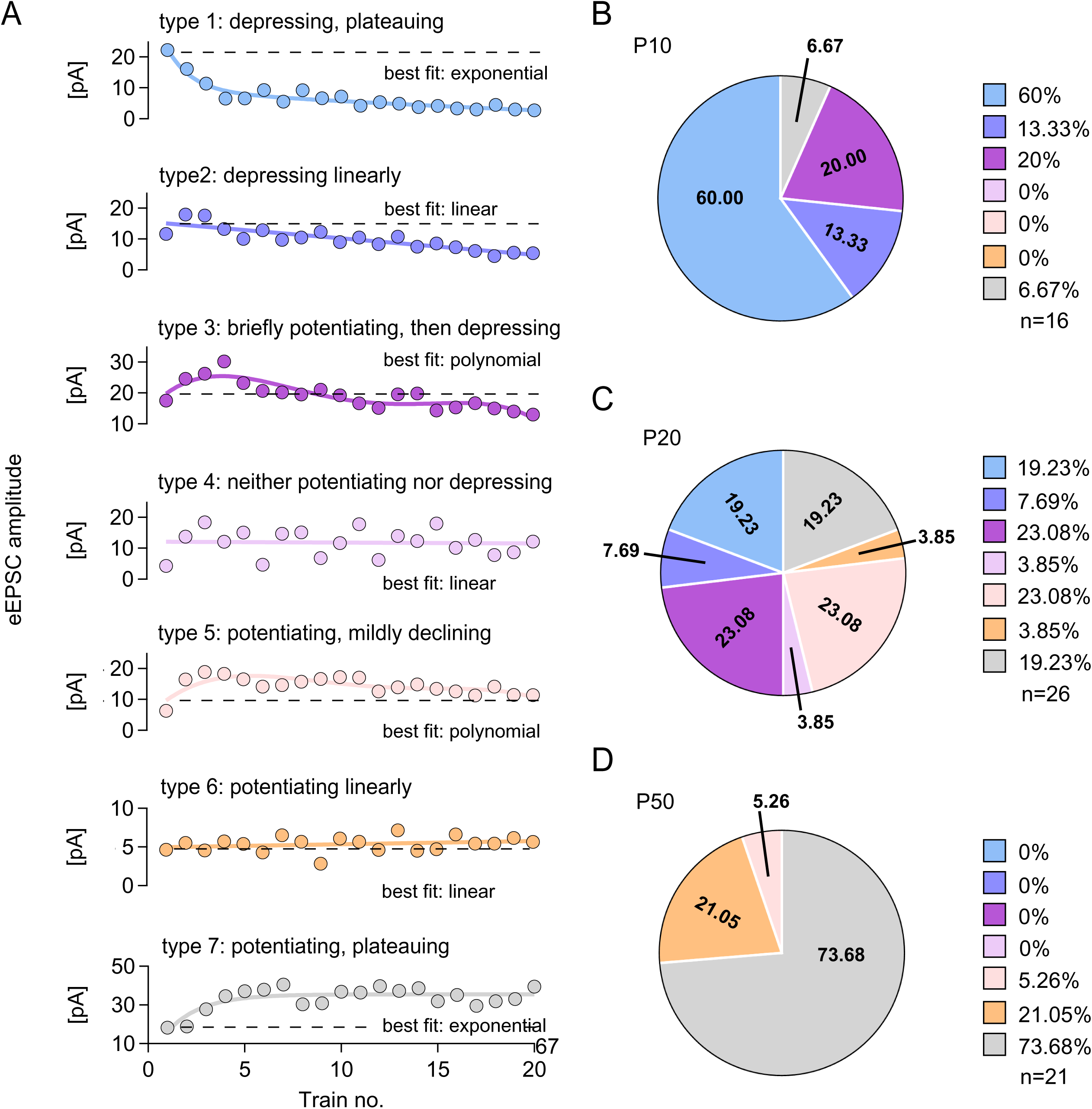
Heterogeneity of STP profiles at neuron-OPC synapses in the corpus callosum. **(A)** : Examples of seven distinct categories based on the shape of the short-term plasticity (STP) profile and the best fit (linear, polynomial, or exponential) during 25 Hz train stimulation. Each example shows the average EPSC amplitude during the phasic (first 5 ms) component of the response after each stimulus in the train, together with the best fit of the data. **(B)** : Pie chart showing the percentage of STP profiles in each of the seven categories in the P10 group, color-coded to match the examples in (A). n = 16 cells from 14 animals. **(C)** : Same as in (B) for the P20 group (n = 26 cells from 21 animals). **(D)** : Same as in (B) for the P50 group (n = 21 cells from 18 animals).

In the P10 group, we observed short-term depression in the majority of OPCs: synapses in 60% of OPCs showed very strong plateauing depression (category 1), 13.33% depressed continuously but without plateauing (category 2), and 20% potentiated briefly before depressing (category 3). A very small group of OPCs (6.67%) displayed a short-term synaptic potentiation (category 7) (Figure 3B). Other time-course patterns were not observed in the P10 group.

In the P20 group, the STP profiles were very heterogeneous between the OPCs. Synapses in 42.31% of OPCs showed short-term depression (19.23% fell into strongly depressing category 1 and 23.08% fell into mainly depressing category 3). Synapses in another 46.16% of OPCs showed short-term potentiation (19.23% fell into strongly potentiating category 7; 23.08% fell into mainly potentiating category 5; 3.85% potentiated linearly (category 6)), (Figure 3C). Synapses in 3.85% of OPCs were neither potentiating nor depressing (category 4).

In the P50 group, we observed short-term potentiation in all OPCs: synapses in 73.68% of OPCs showed strong potentiation (category 7), in 21.05% of cells potentiation was linear (category 6), and in 5.26% of OPCs potentiation occurred with a mild decline (category 5), (Figure 3D).

We found that P10 and P20 were not significantly different (p=0.068; Fisher’s Exact Test), while P10 and P50 groups showed large differences (p=1.00E-06) as did the P20 and P50 groups (p=7.90E-05; Fisher’s Exact Test).

### 2.3. Single-channel conductance of AMPARs in OPCs increases during postnatal development and the increase is likely caused by changes in the subunit composition

Changes in the presynaptic axonal release machinery and/or changes in postsynaptic AMPARs in OPCs may underlie the observed developmental STP dynamics at neuron-OPC synapses. We first analyzed the properties of postsynaptic AMPARs in OPCs. For this, we focused on the quantal current (*q*) which is a basic unit of synaptic transmission, corresponding to the release of one neurotransmitter-filled vesicle (Del Castillo and Katz, 1954; Lisman, 1997). To study *q*, we analyzed delayed EPSCs (dEPSCs) in OPCs which occur after the end of the stimulation train (Figure 2A, 4A). The dEPSCs are considered quantal at neuronal synapses (Korn and Faber, 1991; Redman, 1990), and in our previous study, we reported that dEPSCs have quantal properties also at neuron-OPC synapses (Nagy et al., 2017). The dEPSCs arise due to elevated residual Ca^2+^ at the presynaptic site after the cessation of repetitive stimulation.

We found no significant differences between the age groups in the mean dEPSCs amplitude (p = 0.095; 3.79 ± 0.15 pA at P10; 3.93 ± 0.11 pA at P20, 4.27 ± 0.19 pA at P50), (Figure 4B-C). Rise-time of dEPSCs in P20 group was slightly, but statistically significantly, higher than in P10 and P50 groups (0.521 ± 0.011 ms at P10, 0.570 ± 0.0161 ms at P20, 0.516 ± 0.016 ms at P50; P10 vs P20: p=0.038; P20 vs P50: p=0.038; P10 vs P50: p=0.81), (Figure 4D). Also the decay-time of dEPSCs in P20 group was slightly, but statistically significantly, slower than in P10 and P50 groups (1.292 ± 0.072 at P10, 1.589 ± 0.069 at P20, 1.258 ± 0.075at P50; P10 vs P20: p=0.013; P10 vs P50: p=0.78; P20 vs P50: p=0.0047), (Figure 4E). The total dEPSC charge transfer was slightly higher in P20 than in P10, but not in P50, group (7.485 ± 0.270 fC at P10, 8.533 ± 0.317 fC at P20, 7.928 ± 0.234 fC at P50; P10 vs P20: p=0.043; P10 vs P50: p=0.24; P20 vs P50: p=0.24), (Figure 4F).

**Figure 4:**
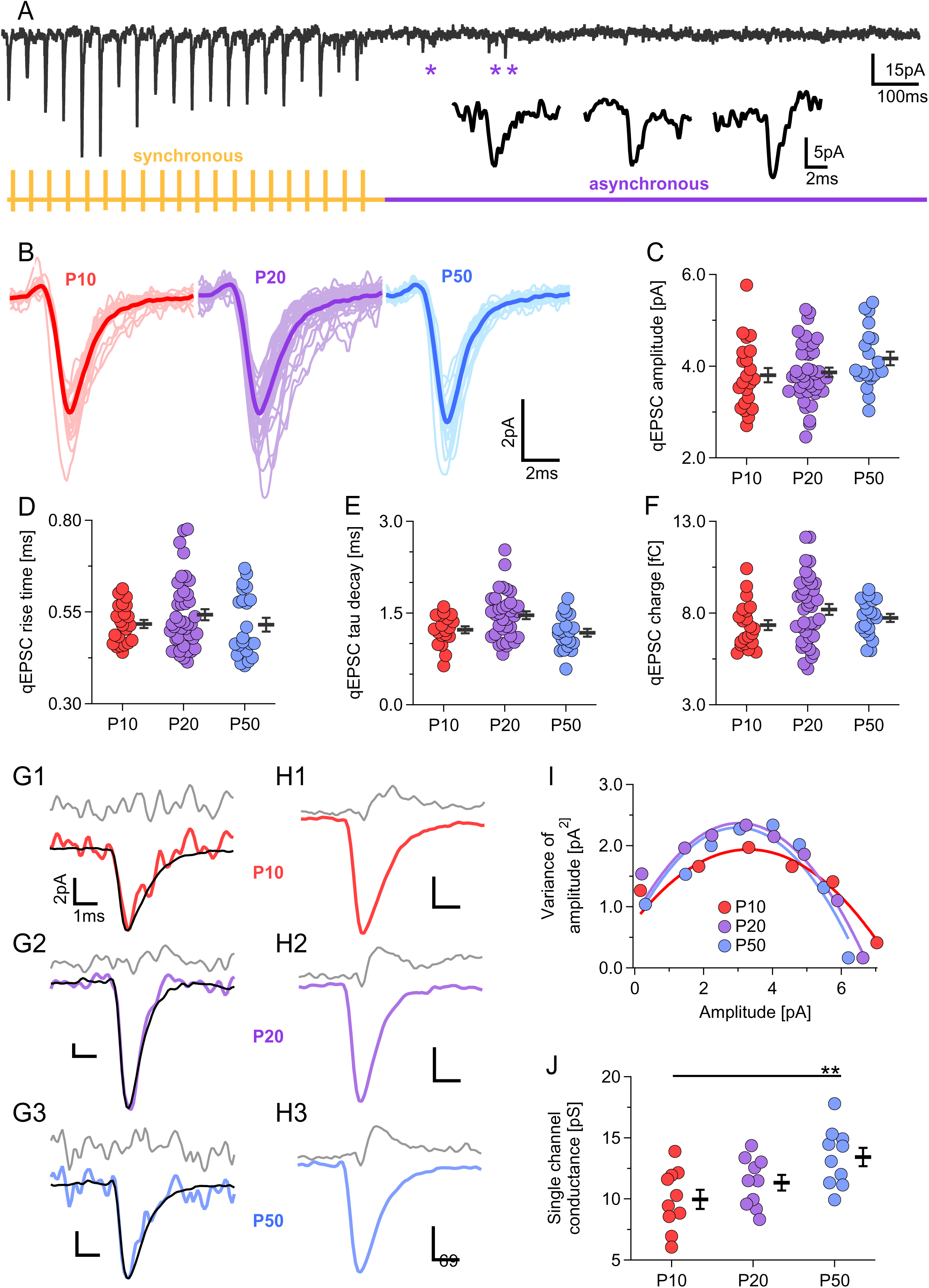
Quantal synaptic currents and single-channel conductance of synaptic AMPARs in OPCs during postnatal development. **(A) :** Top, black: representative trace recorded from a callosal OPC (V_h_ = -80 mV) in response to the train stimulation of the callosal axons, P50 group. Bottom: schematic representation of the stimulation paradigm (20 pulses at 25 Hz) applied to the callosal axons (orange), and a time-period when the delayed EPSCs occur and were collected for the analysis (purple). Asterisks (*) mark examples of individual delayed EPSCs, which are shown on the expanded scale below. Stimulation artifacts are blanked for clarity. **(B)** : Averages of delayed EPSCs from 22 OPCs in P10 group (light red), from 39 OPCs in P20 group (light purple), and from 22 OPCs from the P50 group (light blue). The group means are superimposed on the averages and are shown in dark red (P10 group), dark purple (P20 group), and dark blue (P50 group). **(C)** : Amplitudes of the averaged delayed EPSCs in callosal OPCs. Each circle represents one cell. Black bars represent group average ± SEM. P10 group: n = 25 cells from 22 animals; P20 group: n = 42 cells from 42 animals; P50 group: n = 27 cells from 25 animals. No statistically significant differences were detected between the groups. **(D)** : Rise-time (20-80%) of the delayed EPSCs in callosal OPCs. Each circle represents one cell. Black bars represent group average ± SEM. P10 group: n = 24 cells from 22 animals; P20 group: n = 47 cells from 42 animals; P50 group: n = 27 cells from 25 animals. **(E)** : Decay time of the delayed EPSCs in callosal OPCs. Each circle represents one cell. Black bars represent group average ± SEM. P10 group: n = 22 cells from 22 animals; P20 group: n = 44 cells from 42 animals; P50 group: n = 26 cells from 25 animals. **(F)** : Charge transferred by the delayed EPSCs in callosal OPCs. Each circle represents one cell. Black bars represent group average ± SEM. P10 group: n = 22 cells from 22 animals; P20 group: n = 44 cells from 42 animals; P50 group: n = 24 cells from 24 animals. **(G)** : Non-stationary fluctuation analysis (NSF). Single representative delayed EPSCs in P10 group (red), P20 group (purple), P50 group (blue), overlaid with the re-scaled average EPSC waveform (black) recorded from the same cells. Grey trace represents the residual noise for the corresponding event (i.e. after the subtraction of the black trace from the corresponding colored trace). **(H)** : Average EPSCs waveforms from a representative random event selection in the P10 group (red), P20 group (purple), and P50 group (blue). The gray traces represent the mean residual noise variance across all delayed EPSCs for the corresponding random event selection. **(I)** : Mean-variance plot of the amplitude of the delayed EPSCs in OPCs obtained during NSF, and the corresponding parabola fits in the P10 group (red), P20 group (purple), and P50 group (blue). **(J)** : Single channel conductance of AMPARs in OPCs obtained after 10 repetitions of random events selection within each age-group. Each colored dot represents a single random measurement. Black bars represent mean ± SEM. ** = p<0.01.

To study single-channel conductance of synaptic AMPARs in OPCs, we performed non-stationary fluctuation analysis (NSFA). We found that single-channel conductance changed significantly during animal maturation: 9.96 ± 0.78 pS in P10 group, 11.33 ± 0.64 pS in P20 group, and 13.42 ± 0.75 pS in P50 group (Figure 4G-J).

Single-channel conductance of AMPARs varies depending on their subunit composition (Cull-Candy et al., 2006; Greger et al., 2017; Traynelis et al., 2010): GluA2-lacking AMPARs have higher single-channel conductance in neurons and in OPCs (Chen et al., 2018; Liu and Zukin, 2007; Swanson et al., 1997). Hence, the observed difference between the age-groups in the single-channel conductance of synaptic AMPARs may be associated with subunit composition or changes in the presence or editing of the GluA2 subunit within the AMPARs complex during animal maturation (Coombs and Cull-Candy, 2021). To investigate this, we studied the current-voltage (I-V) relationship of AMPAR-mediated EPSCs in callosal OPCs in the presence of spermine in the pipette solution, and analyzed the rectification index (RI) (Figure 5A-B). We found that in P10 mice, the I-V curves were close to linear (RI = 0.364 ± 0.020), while in older mice they were inwardly rectifying: RI=0.157 ± 0.030in P20 mice, and RI = 0.138 ± 0.030 in P50 animals. Statistical analysis showed difference between P10 and P20, and P10 and P50 groups: P10 vs P20, p= 3.18E-05; P10 vs P50, p= 2.42E-05, while P20 and P50 groups did not test as different (p=0.627) (Figure 5A-B). Thus, the proportion of edited GluA2 subunit within AMPAR complex is lower in OPCs from more mature vs very young mice, and synaptic AMPARs become more Ca^2+^-permeable with animal maturation. To explore this issue further, we adopted a model described by Stubblefield and Benke (Stubblefield and Benke, 2010), which translates rectification index (RI) to the relative proportions of Ca^2+^-permeable to Ca^2+^-impermeable AMPARs (Figure 5C-D). Based on this model, we found that only 19.59 ± 4.99% of the synaptic AMPARs were Ca^2+^-permeable in the P10 group, their proportion increased to 68.26 ± 6.24% in P20 animals, and further increased to 73.66 ± 6.79 % in P50 mice (P10 vs P20, p=1.24E-05; P10 vs P50, p=6.89E-06), (Figure 5C-D).

**Figure 5:**
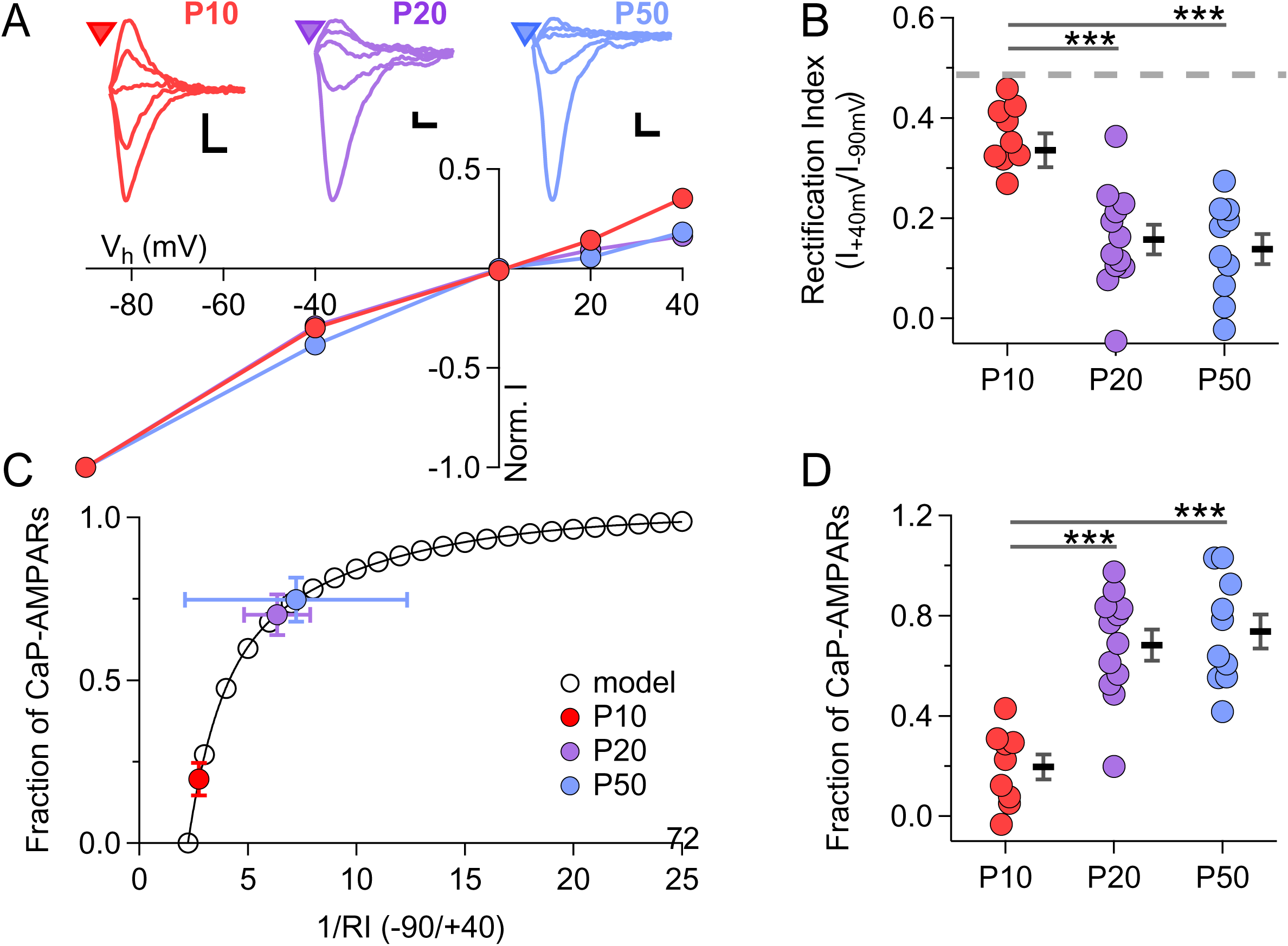
AMPARs in OPCs become more Ca^2+^ - permeable during callosal development. **(A)**: Current-voltage (I-V) relationship for the phasic EPSCs recorded in OPCs in three age groups. Each dot represents the mean EPSC amplitude ± SEM in the age group, recorded at V_h_ = -90, -40, 0, +20 or +40 mV, and normalized to the average amplitude recorded at V_h_ = -90 mV. Examples of the averaged EPSCs are shown above the I-V plot. P10 group (red): n = 9 OPCs from 6 animals; P20 group (purple): n = 12 OPCs from 9 animals; P50 group (blue): n = 10 OPCs from 9 animals. **(B)** : Summary graph showing the rectification index (RI) for the cells shown in (A). Each colored dot represents the RI of one OPC. Black bars represent group mean ± SEM. Grey dashed line indicates the theoretical perfect, non-rectifying I-V relationship (RI = 0.44). * = p<0.05; ** = p<0.01; *** = p<0.001. The number of cells and animals is as in **(A)**. **(C)** : Model relationship between the inverted rectification index (1/RI) and fraction of the rectifying, Ca^2+^-permeable AMPARs (CP-AMPARs). Colored dots represent average 1/RI for each experimental group plotted against the mean calculated CP-AMPAR fraction ± SEM. The number of cells and animals is as in **(A)**. **(D)** : Summary of the fraction of CP-AMPARs in all age groups, based on **(C)**. Each colored dot represents the CP-AMPAR fraction in one cell. Black bars represent group mean ± SEM. * = p<0.05; ** = p<0.01; *** = p<0.001. The number of cells and animals is as in **(A)**.

Considering that Ca^2+^-permeable AMPARs play a role in the mechanisms of STP at neuronal synapses (Rozov et al., 2018), our findings suggest that postsynaptic mechanisms may contribute to the switch from short-term depression to short-term facilitation which we observed upon animal maturation (Figure 2).

### 2.4. Computational modelling suggests that filtering effects underlie differences in quantal EPSC amplitude at synaptic sites of neuron-OPC synapses vs OPC cell soma

We were curious why the higher single-channel conductance of AMPARs in older vs younger mice and an increased ratio of Ca^2+^-permeable AMPARs did not result in higher quantal EPSC amplitudes in the older mice. The relationship between synaptic current amplitude (I_EPSC_) and single-channel conductance (*γ*) is determined by the formula:

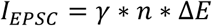

where *n* is the number of AMPAR channels, and Δ*E* is the driving force. As the driving force remained constant in our experiments, a possible explanation of our finding is that the number of activated AMPARs decreases in older animals. An alternative explanation might be that lower membrane resistance (Figure 1A), more complex cell morphology of older OPCs (Kula and Kukley, 2025), and the corresponding in increase in C_m_ (Figure 1C) contribute to higher electrotonic filtering (Spruston et al., 1994; Williams and Mitchell, 2008). Therefore, it is possible that the local synaptic conductance at distal processes, where the synaptic transmission happens, is larger in OPCs from older animals, but EPSCs at the cell soma remain similar due to the stronger filtering effect while traveling from the local site to the cell soma. We employed a computational modeling approach to investigate which of the two theories is more likely. To consider the complex highly ramified morphology of OPC, in our modelling we used the morphological parameters of those OPCs which we have analyzed in detail and described previously (Kula and Kukley, 2025). We created passive OPC models using the reconstructed individual OPC morphology data (n=12 cells for each age group). We set the diameter of all OPC processes to be 0.2 µm and the diameter of the OPC soma to be 6.5 µm, similar to previously reported (Sun and Dietrich, 2013).

In the modelling, we considered the OPC’s response to a single synaptic input and used a model for the synaptic current in the form of *I*_*syn*_ = *g*_*syn*_*s*(*t*)(*V* − *E*_*syn*_). The model was implemented using NetStim within NEURON with *E*_*syn*_ = 0 *mV* and *s*(*t*) a double exponential with rise time *τ*_1_ = .25 ms and decay time *τ*_2_ = 1 ms.

In the first set of simulations, we assumed that the number of AMPAR channels remained constant across P10, P20 and P50 groups. Therefore, the local synaptic conductance is proportional to single AMPAR channel conductance. Let 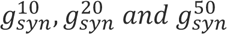 denote the local synaptic conductance of cells within the p10, p20 and p50 age groups, respectively. Following (Sun and Dietrich, 2013), we assume that 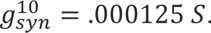 We then choose 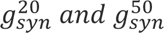 so that the ratios 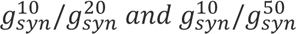 are the same as that for the corresponding single AMRAR conductances. This gives 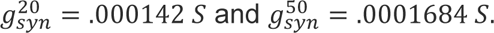

We stimulated each model cell within each age group at a randomly chosen location and determined the corresponding maximum somatic current 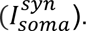 First, we performed one iteration of simulations from each model cell within its corresponding age group, N = 12 model cells in P10, P20 and P50 groups), at a randomly chosen location. Statistical analysis showed that the maximum somatic current is comparable (p > 0.05) across each pair of age groups (Figure 6A). Subsequently, we ran 100 iterations of the simulations with different randomly chosen synaptic locations in each OPC model and computed p-values comparing peak 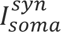 across the age groups, for each iteration (Figure 6B). Over 75% of the iterations generated comparable peak 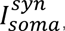 consistent with what we observed with experimental data (Figure 4), despite the local synaptic conductance increasing from P10 to P50.

**Figure 6:**
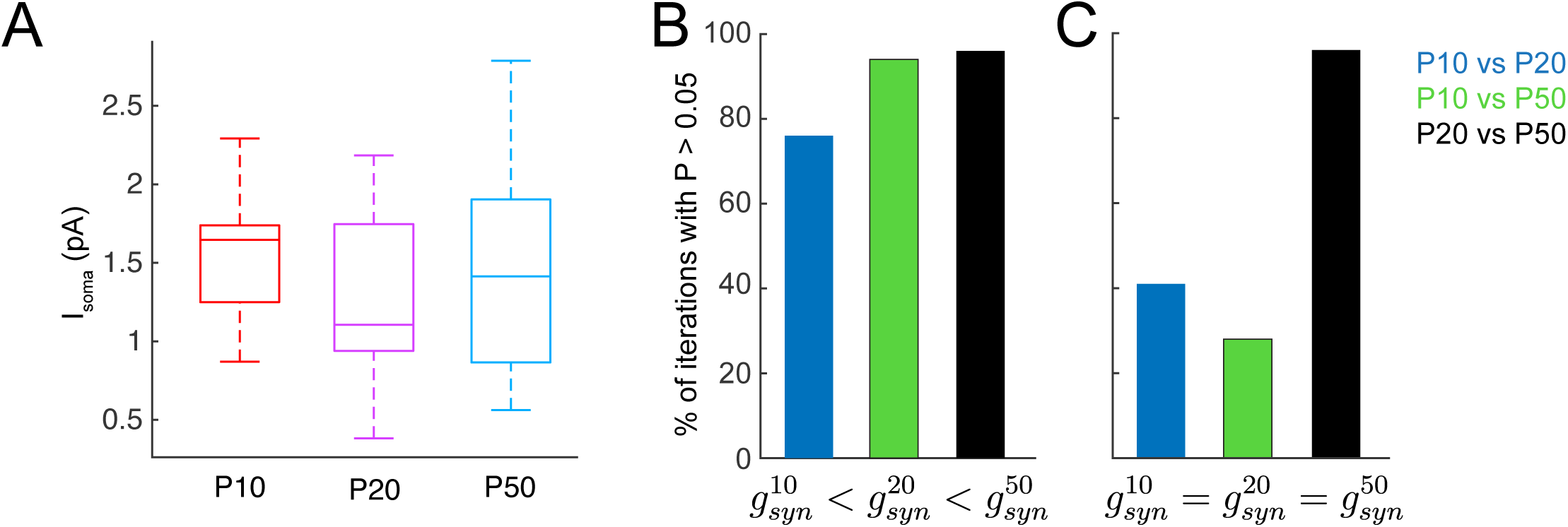
Computational modelling suggests that filtering effects underlie differences in quantal EPSC amplitude at synaptic sites of neuron-OPC synapses vs OPC cell soma. Passive OPC models were created using reconstructed individual OPC morphology data (Kula and Kukley, 2025) (n = 12 cells each for P10, P20 and P50 groups). A single excitatory synaptic event was simulated with double-exponential kinetics defined by τ_rise_ and τ_decay_. **(A)** : A single excitatory synapse was randomly placed on each passive OPC model morphology tree and the somatic current (I_soma_) was simulated. The synaptic conductance for P10, P20 and P50 groups were set as following: 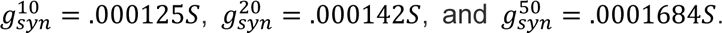 Statistical analysis showed that the maximum somatic currents were comparable across each pair of age groups (p > 0.05 for all pair-wise multi-group comparison, student t test). **(B)** : The same type of simulation as in **A** was repeated for 100 iterations with the synapses being randomly placed at different positions. For each iteration, p values comparing peak somatic currents across three age groups were obtained. The Bar graph shows the percentage of iterations that demonstrates p value > 0.05. Our simulation shows that the majority of iterations show comparable somatic current amplitudes across age groups, consistent with our experimental data. **(C)** : In a separate set of simulation, we set identical synaptic conductance for all three age groups 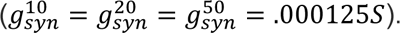 We ran a different set of 100 iterations and obtained p values comparing somatic current amplitudes across age groups. The simulation demonstrates a substantial proportion of iterations with significant differences between the P10 and P20 or P50 groups, contradicting our experimental data.

In the second set of simulations, we assumed that the number of single AMPAR channels decreases in older animals so that the local *g*_*syn*_ remains comparable in all three age groups (as the number of channels decreases but the single channel conductance increases). We assumed that 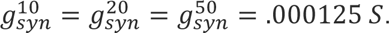 We then ran another 100 iterations as we did above and calculated the numbers of iterations that show comparable peak 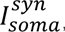 (p>0.05) among pairs of age groups. We found a difference of the 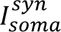 between P10 and P20 groups of animals as well as between and P10 and P50 mice (Figure 6C). This result of the modelling did not reflect our experimental data. (Note that little difference in 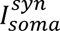 between P20-P50 groups which we observed in our modeling may be related to the fact that OPCs in P20 and P50 cells are roughly the same size and larger than OPCs in P10 animals).

Taken together, the results of our computational modeling support the theory that the single-channel conductance of synaptic AMPARs is larger in OPCs from older than from younger mice, while the numbers of single AMPAR channels is comparable. This results in a larger local synaptic conductance, and hence in larger quantal current amplitude at neuron-OPC synapses in older vs younger mice. However, when the synaptic current travels from synapses to the OPC cell soma, it undergoes a stronger filtering effect in OPCs of older mice than in OPCs of younger mice due to the increased morphological complexity in their cellular process architecture. Hence, the quantal currents at the cell soma appear comparable between the age-groups.

### 2.5. Neuron-OPC synapses have low release probability, and its dynamic during stimulation trains follows the dynamics of the synaptic strength

In the previous part, we focused on the postsynaptic sites of neuron-OPC synapses. Here, we switched to the analysis of maturation-dependent changes at the presynaptic (axonal) sites of these synapses.

First, we estimated the basal release probability which is the likelihood that a presynaptic action potential will trigger a fusion of at least one synaptic vesicle filled with neurotransmitter at a given release site. It is usually considered that the response probability reflects the release probability at synapses. For these experiments, we used a minimal stimulation paradigm (see Materials and Methods) (Figure 7A-B). The response probability was almost identical in all three investigated age-groups: 0.0975 ± 0.0140 at P10, 0.0956 ± 0.0152 at P20, 0.0737 ± 0.0186 at P50; and no statistically significant differences were observed (P10 vs P20: p=0.93; P10 vs P50: p=0.73; P20 vs P50: p=0.73), (Figure 7C). Thus, the basal release probability at neuron-OPC synapses in the corpus callosum is low, and does not change during postnatal development.

**Figure 7:**
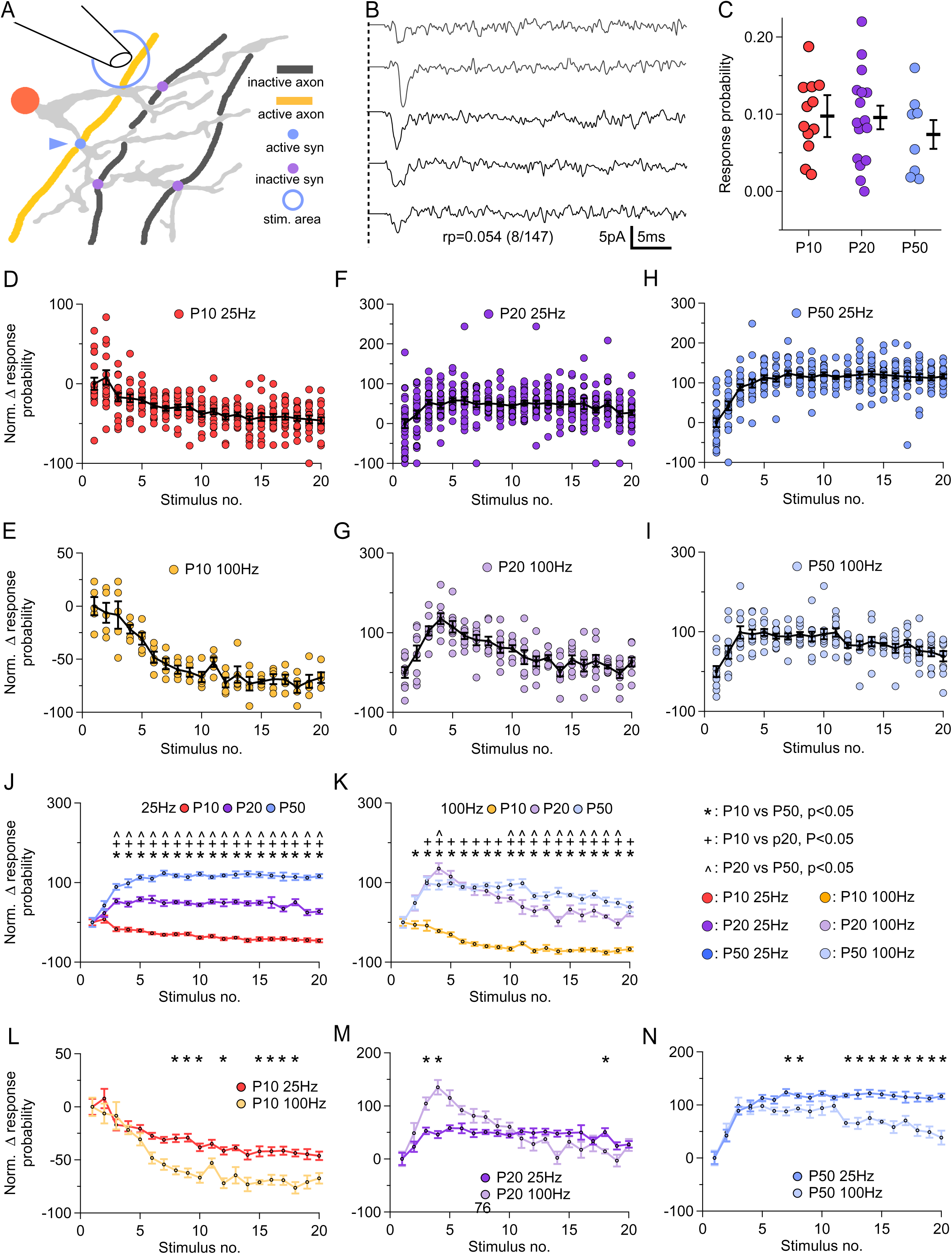
Neuron-OPC synapses have low release probability, and its dynamic during stimulation trains follows the dynamics of the synaptic strength. **(A)** : Schematic drawing showing the minimal stimulation paradigm. An OPC (shown with orange cell soma and grey cell processes) has synaptic contacts with 3 axons (one shown in yellow and two shown in black; synaptic contacts are shown with blue and purple dots). During the minimal stimulation (low voltage), only the yellow axon is activated and releases synaptic vesicles at its synaptic contacts (indicated by a blue dot) with the OPC. The black axons are not recruited by the stimulation and do not release synaptic vesicles despite having synaptic contacts with the OPC. **(B)** : Representative example traces showing EPSCs in OPCs (P50 group) in response to the minimal stimulation of callosal axons. The dashed line represents the beginning of the stimulation. Stimulation artifacts are removed for clarity. **(C)** : Probability of recording an EPSC in response to the first stimulus of the minimal stimulation train of 20 stimuli at 25 Hz in three age-groups. Each circle represents one cell. Black bars represent group average ± SEM. P10 (red): n = 12 cells from 11 animals; P20 group (purple): n = 16 cells from 15 animals; P50 group (blue): n = 8 cells from 8 animals. None of the groups tested as significantly different (One-way ANOVA, p=0.593). **(D)** : Dynamics of the response probability at neuron-OPC synapses during the train. Each circle represents a normalized average probability of recording an EPSC in a single OPC during the train of 20 pulses at 25 Hz. Black line with circles represents the group mean ± SEM; n = 15 cells from 14 animals. **(E)** : As **(D)**, but for the train stimulation of the callosal axons with 20 pulses at 100 Hz; n = 14 cells from 10 animals. **(F)** : As **(D)**, but for the P20 group; n = 23 cells from 18 animals. **(G)** : As **(E)**, but for the train stimulation of the callosal axons with 20 pulses at 100 Hz; n = 19 cells from 14 animals. **(H)** : As **(D)**, but for the P50 group; n = 19 cells in 16 animals. **(I)** : As **(E)**, but for the train stimulation of the callosal axons with 20 pulses at 100 Hz; n = 9 cells from 9 animals. **(J)** : Summary of **(D)**, **(F)** and **(H)**: probability to record an EPSC in an OPC upon stimulation of callosal axons with 20 stimuli at 25 Hz in three age-groups. Each circle represents the group mean ± SEM. P10 in red, P20 in purple, P50 in blue.* = p<0.05, P10 vs P50. ^+^ = p<0.05, P10 vs P20. ^#^ = p<0.05, P20 vs P50. **(K)** : Summary of **(E), (G) and (I)**: probability to record an EPSC in an OPC upon stimulation of callosal axons with 20 stimuli at 25 Hz in three age-groups. Each circle represents the group mean ± SEM. P10 in orange, P20 in purple, P50 in blue.* = p<0.05, P10 vs P50. ^+^ = p<0.05, P10 vs P20. ^#^ = p<0.05, P20 vs P50. **(L)** : Summary of **(D)** and **(E)**: Comparison of the response probabilities during 20 stimuli train at 25 Hz vs 100 Hz, for the P10 group. * = p<0.05. **(M)** : Summary of **(F)** and **(G)**: Comparison of the response probabilities during 20 stimuli at 25 Hz vs 100 Hz, for the P20 group. * = p<0.05. **(N)** : Summary of **(H)** and **(I)**: Comparison of the response probabilities during 20 stimuli at 25 Hz vs 100 Hz, for the P50 group. * = p<0.05.

Next, we analyzed the response probability during stimulation trains. In the P10 group, the response probability was decreasing throughout the whole 25 Hz train, reaching a reduction by - 46.16 ± 4.01% by the end of the train (Figure 7D). The decrease was almost twice more pronounced during the 100 Hz train with the response probability being reduced by -67.36 ± 4.92% at the end of the train (Figure 7E). In the P20 group, the response probability was rising during the first 3-5 pulses, by +58.03 ± 5.29% and +134.98 ± 13.59% at frequencies of 25 or 100 Hz, respectively (Figure 7F-G). During the subsequent pulses, the response probability was decreasing although still remained potentiated, and by the end of the train it was still higher than the initial value by +27.34 ± 6.12% and +26.34 ± 11.33% for 25 and 100 Hz stimulation, respectively (Figure 7F-G). In the P50 group, upon 25 Hz stimulation, the response probability was increasing during the first 10 stimuli and remained stable till the end of the train reaching +116.14 ± 5.13% of the initial value by the end of the train (Figure 7H). Upon 100 Hz stimulation, the response probability increased during the first three stimuli reaching +98.22 ± 15.71% of the initial value but decreased during the subsequent pulses, ending at +38.51 ± 13.04% higher value than the initial one (Figure 7I).

Statistically, the dynamic profiles of response probability during the trains were significantly different between the groups (Figure 7J-K). For 25 Hz stimulation, significant differences were observed starting at stimulus 3 (P10 vs P20: p<1E-15 to p=5.94E-07; P10 vs P50: p<1E-15 to p=1.45E-10; P20 vs P50: p<1.00E-15 to p=4.98E-04), (Figure 7J). For 100 Hz stimulation, we found significant differences between the P10 and P20 or P50 groups almost throughout the entirety of the stimulation (P10 vs P20: p=2.28E-06 to p=0.044; P10 vs P50: p=3.20E-14 to p=0.037) paradigms (Figure 7K). Consistent statistically significant differences between P20 and P50 groups were observed starting from the tenth stimulus in the train (p=0.0058 to p=0.044) (Figure 7K). We also performed statistical comparison of the response probability values between the two stimulation frequencies within each age-group (Figure 7L-N). We found mostly consistent differences between the stimulation paradigms in the P10 and P50 groups: 3.30E-7 to p=0.0030 at P10, p = 1.87E-6 to p=0.013 at P50, but the P20 group only showed consistent differences at stimuli 3 and 4.

When comparing together the time-course of the response probability (Figure 7D-I) and the time-course of the EPSC amplitudes (Figure 2A-F), we found that they showed very similar dynamics (although not absolutely identical). As response probability at synapses is directly related to the presynaptic release probability, our findings suggest that presynaptic mechanisms play an important role for the observed dynamic changes of STP at neuron-OPC synapses.

### 2.6. Number of vesicles that can be released at neuron-OPC synapses during stimulation of callosal axons does not change during postnatal development

As neuron-OPC synapses in P10 animals exhibited short-term depression whereas in older animals they showed facilitation, we asked whether developmental differences in the synaptic vesicle output could contribute to these distinct plasticity profiles. To estimate the number of vesicles which can be released at neuron-OPC synapses in our experimental conditions, we divided the charge transferred by the phasic EPSC evoked during maximal stimulation by the charge transferred during quantal EPSCs. Maximal EPSCs were obtained using a strong stimulation that allows activating all callosal axons that can be recruited in our stimulation conditions (see Materials and Methods; (Kukley et al., 2007)) Figure 8A-C).

**Figure 8:**
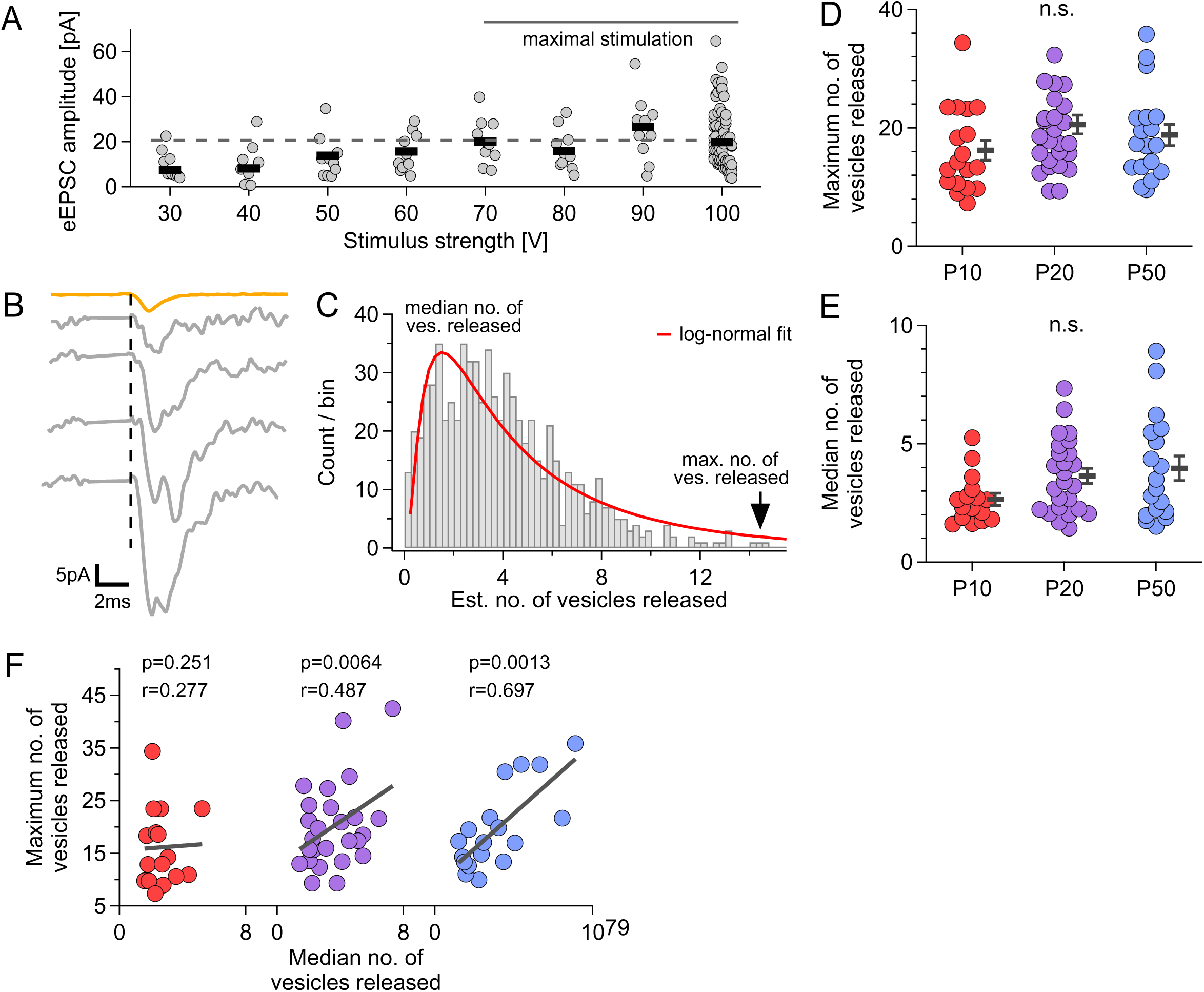
The number of vesicles released at neuron-OPC synapses does not differ between the age-groups. **(A)** : Amplitudes of EPSCs evoked in a single P20 OPC by stimulation of increasing strength, from 30 V to 100 V, in +10 V increments. Each circle represents the amplitude of a single phasic EPSC; black bars represent the mean amplitude at each stimulation strength. The dashed gray line represents the maximal average amplitude reached at the highest stimulation intensities (70-100 V) and reflects activation of callosal axons that can be recruited by our stimulation. **(B)** : Examples of phasic EPSCs recorded during maximal stimulation (grey) and an averaged quantal qEPSC (orange) from the same cell, all shown with the same scaling. The black dashed line represents stimulus artifacts (blanked for clarity). **(C)** : Histogram distribution of the ratios between charges transferred during maximal EPSC and qEPSC from the cell shown in (A). This ratio was used to estimate the number of neurotransmitter vesicles released during stimulation. The red line represents the best log-normal fit to the distribution. The black arrow indicates the largest number of vesicles released during stimulation (∼15 vesicles). **(D)** : The maximum number of vesicles released during maximal stimulation compared between the age groups. Each circle represents the maximum value from a single cell; black bars represent the group mean ± SEM. P10 in red, n=18 cells in 17 animals; P20 in purple, n=28 cells in 27 animals; P50 in blue, n=18 cells in 16 animals. **(E)** : The median number of vesicles released during maximal stimulation compared between age groups. Each circle represents the median value from a single cell; black bars represent the group mean ± SEM. P10 in red, n=16 cells in 16 animals; P20 in purple, n=25 cells in 25 animals; P50 in blue, n=18 cells in 16 animals. **(F)** : The maximum number of vesicles released plotted against the median number of vesicles released. Each circle represents a single cell. Black lines represent a linear fit of the data.

We found no significant differences between the age groups in either the maximal number of vesicles released (16.35 ± 1.68 vesicles in the P10 group, 20.85 ± 1.57 vesicles in the P20 group, and 19.63 ± 1.86 vesicles in the P50 group; Kruskal-Wallis test, p = 0.138) or in the median number of vesicles released (2.67 ± 0.25 vesicles in the P10 group, 3.58 ± 0.31 vesicles in the P20 group, and 3.94 ± 0.52 vesicles in the P50 group; Brown-Forsythe ANOVA test, p = 0.086; Figure 8D-E). These findings suggest that differences in vesicle availability are unlikely to account for the short-term depression observed in P10 animals compared with facilitation in older mice.

Finally, when we correlated the maximal and median numbers of released vesicles, we found a moderate positive correlation in the P20 group (Spearman’s r = 0.487; p = 0.0064) and a strong positive correlation in the P50 group (r = 0.697; p = 0.0013), but no significant correlation in the P10 group (r = 0.277; p = 0.251; Figure 8F). This suggests that the relationship between maximal vesicle output and typical release becomes stronger during maturation, indicating increased consistency of synaptic release at the synapses.

### 2.7. Asynchronous release significantly contributes to the total glutamate release at neuron-OPC synapses during the train

To investigate dynamic changes at presynaptic and postsynaptic sites of neuron-OPC synapses, we have so far analyzed phasic EPSCs and dEPSCs. But besides those two types of synaptic currents, we have also observed EPSCs of small amplitude triggered after each stimulus during the train. To analyze those asynchronous EPSCs (aEPSCs) (Figure 9A), we studied their count after each pulse during the 25 Hz trains and their amplitudes. For this analysis, we considered that aEPSCs are the synaptic currents with an onset of >5 ms after each stimulus. (We have not analyzed aEPSCs during 100 Hz trains because the interval between the sequential pulses in those trains is only 10 ms and, assuming that the first 5 ms correspond to the phasic response, the remaining time of 5 ms is too short to reliably sample the aEPSCs after each pulse).

**Figure 9:**
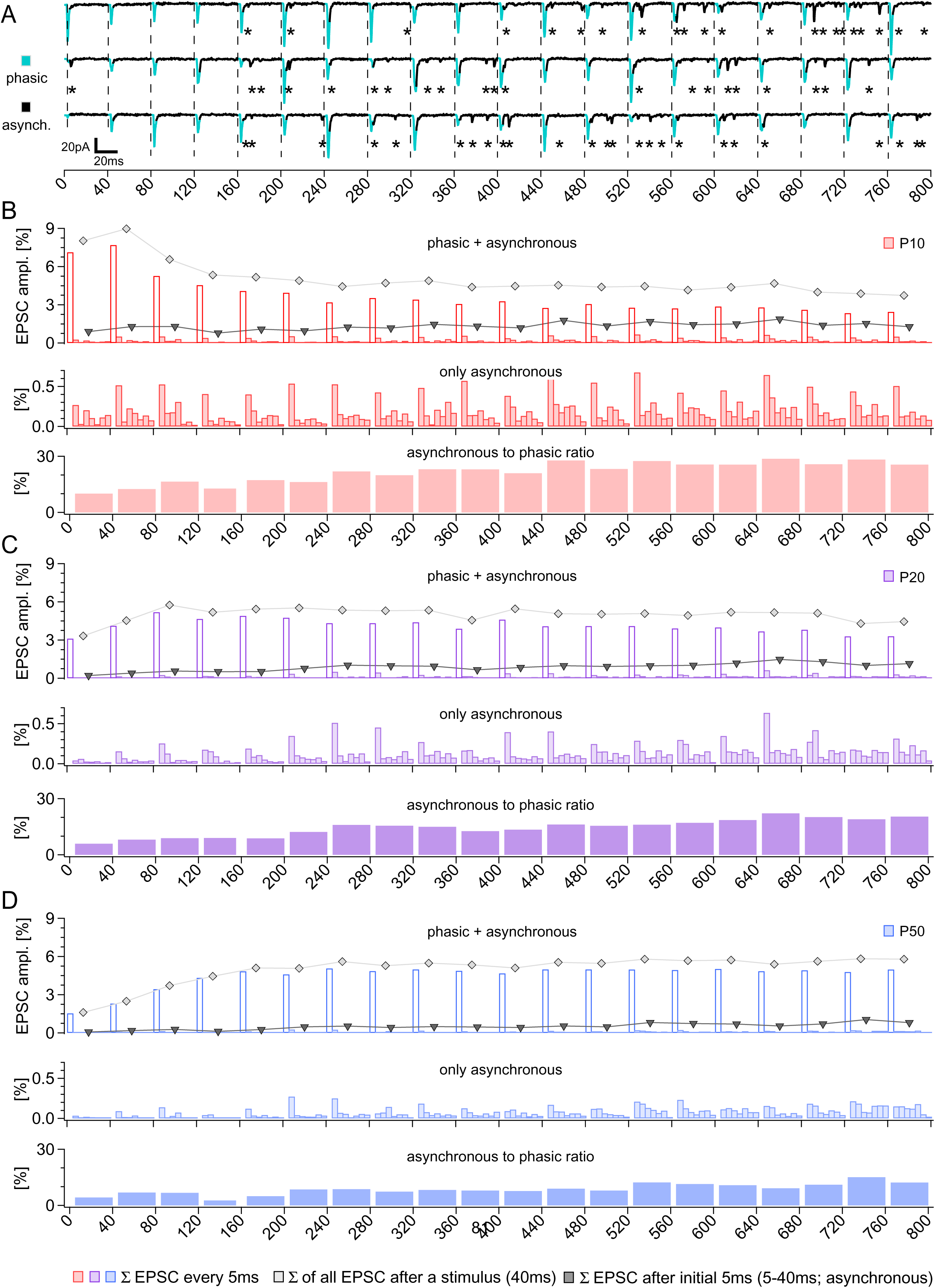
Asynchronous release significantly contributes to the total glutamate release at neuron-OPC synapses during the train. **(A) :** Three example traces showing phasic and asynchronous EPSCs evoked in an OPC (from P50 group) by a train of 20 stimuli at 25 Hz. Phasic EPSCs are marked by turquoise color. Asynchronous EPSCs are marked by asterisks (*). **(B) :** *Top panel:* The mean fraction (%) of the EPSCs evoked by a train (20 stimuli at 25 Hz), binned every 5ms, in the P10 group. Each colored bar represents the ratio of EPSCs (phasic + asynchronous) count within each 5 ms bin to the total number of EPSCs within the whole train. The line with gray diamonds indicates the summed percentage of all (phasic + asynchronous) EPSCs triggered after a given stimulus during the train. The line with black triangles indicates the summed percentage of the asynchronous EPSCs (i.e. the EPSCs which fulfil the following condition: 5 ms < onset time ≤40 ms) triggered after a given stimulus during the train. *Middle panel:* The mean fraction (%) of the asynchronous EPSCs evoked by a train (20 stimuli at 25 Hz), binned every 5ms, in the P10 group. Each colored bar represents the ratio of asynchronous EPSCs within each 5 ms bin to the total number of EPSCs within the whole train. *Bottom panel:* Each colored bar represents the ratio between the asynchronous and the phasic EPSCs count after each stimulus during the train, for the P10 group. **(C) :** As **(B)** but for the P20 group. **(D) :** As **(B)** but for the P50 group.

To investigate how the occurrence of asynchronous release is related to the total release during the train, we separated phasic EPSCs and aEPSCs, and binned aEPSCs into the consecutive 5 ms categories (see Materials and Methods). We then calculated the ratio between the charge transferred during each 5 ms bin and the total charge within a train. After the first stimulus, aEPSCs contributed 10.04%, 5.93%, and 4.27% to the total release during the train in P10, P20, and P50 groups, respectively (Figure 9B-D). However, their proportion continuously increased with successive stimuli during the train, reaching 25.55% at P10, 20.47% at P20 and 12.24% at P50 by the end of the train (Figure 9B-D). Hence, our data suggest that asynchronous release makes a significant contribution to the total neurotransmitter release during train stimulation, and its contribution increases during the time-course of the train.

### 2.8. Changes in the synaptic latency at neuron-OPC synapses during train stimulation are larger in the juvenile than in the adult animals

In neuronal networks, an important parameter that influences information transmission is synaptic latency which is defined as the time-interval between a presynaptic action potential and the postsynaptic response. Two major factors determine the synaptic latency: the axonal conduction time and the synaptic delay (Boudkkazi et al., 2011; Sabatini and Regehr, 1999). Conduction time refers to the time it takes for an action potential to travel along the presynaptic axon and reach the synaptic bouton. Synaptic delay refers to the time which is required for neurotransmitter release, diffusion across the synaptic cleft, and receptor binding in the postsynaptic cell. Synaptic latency is crucial for efficient and coordinated neural communication, and variations in synaptic latency can influence the timing and the strength of synaptic transmission, impacting neuronal firing and circuit dynamics. In analogy, at neuron-OPC synapses synaptic latency is likely to influence the coordination of fast axon-glia transmission and communication within neuron-glia circuits. Hence, we studied changes in the synaptic latency for the phasic EPSC at axon-OPC synapses during train stimulation (Figure 10A-B).

**Figure 10:**
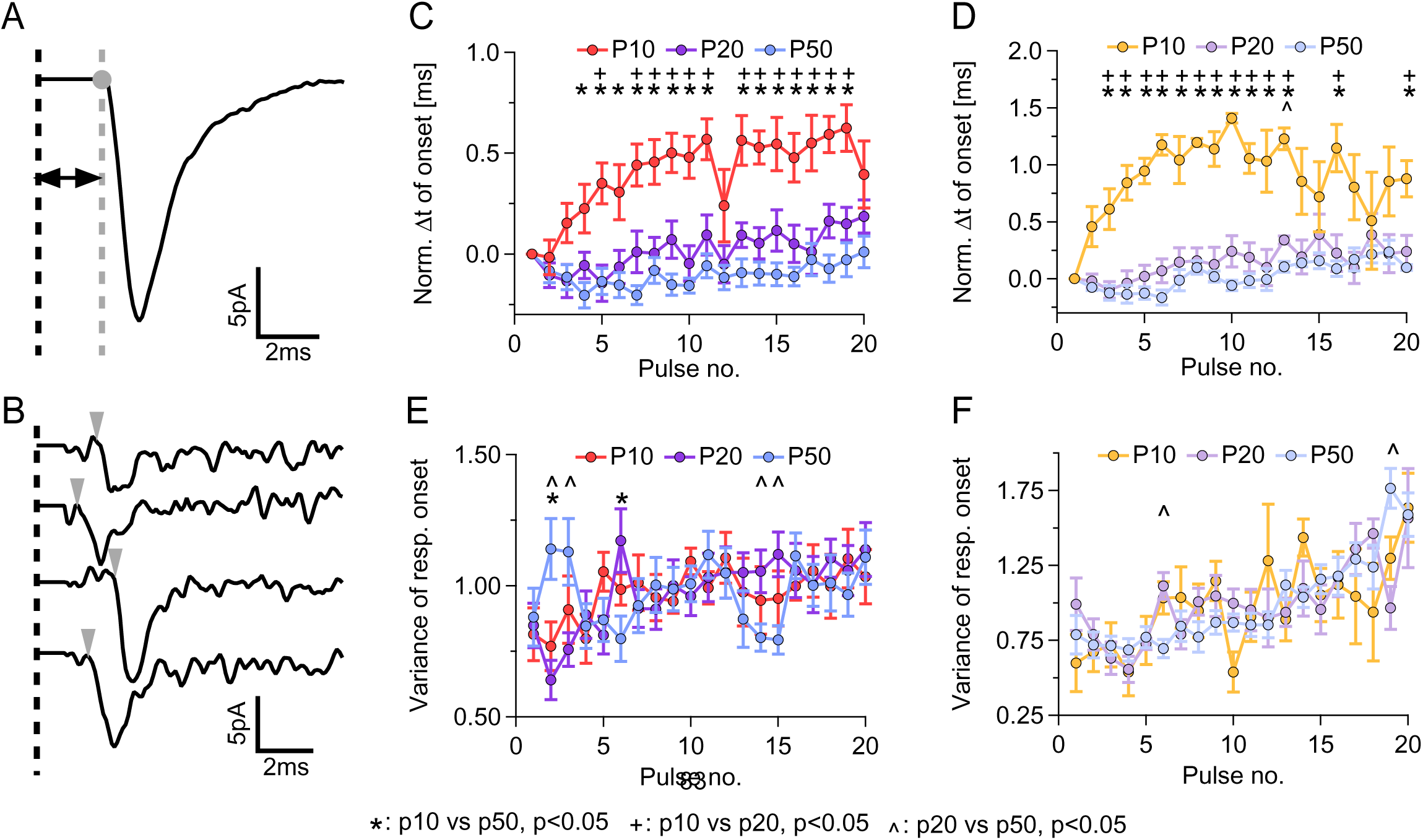
Changes in the synaptic latency at neuron-OPC synapses during train stimulation are larger in the juvenile than in the adult animals. **(A) :** Example of an averaged phasic EPSC recorded in an OPC in response to the first stimulus in the train (P50 group). The black dashed line indicates the start of the stimulation (the stimulation artifact is removed for clarity). The grey dashed line and circle represent the onset of the phasic EPSC. Black arrows mark the synaptic latency, i.e. the time between the start of the stimulus and the onset of the phasic EPSC. **(B) :** Examples of individual phasic EPSCs with various onsets triggered by the first stimulus in the train. The EPSCs were recorded in the same OPC during 4 trials of the callosal axons stimulation. The EPSCs were isolated and plotted together to highlight the variability in their synaptic latencies. The black dashed line indicates the start of the stimulation (the stimulation artifact is removed for clarity). The gray arrowheads indicate the time points of the EPSC onsets. **(C) :** Dynamics of the synaptic latencies of the phasic EPSCs recorded during the trains of 20 pulses at 25 Hz in OPCs from three age groups. Each circle represents the normalized group mean latency after a given stimulus, mean ± SEM. P10 group (red): n = 15 cells from 14 animals; P20 group (purple): n = 23 cells from 18 animals; P50 group (blue): n = 19 cells from 16 animals. * = p<0.05, P10 vs P50; ^+^ = p<0.05, P10 vs P20; ^#^ = p<0.05, P20 vs P50. **(D) :** As **(C)** but for trains of 20 pulses at 100 Hz. P10 group (orange): n = 14 cells from 10 animals; P20 group (purple): n = 19 cells from 14 animals; P50 group (blue): n = 9 cells from 9 animals. * = p<0.05, P10 vs P50; ^+^ = p<0.05, P10 vs P20; ^#^ = p<0.05, P20 vs P50. **(E) :** Dynamics of the variance of the synaptic latency of phasic EPSCs during the train of 20 stimuli at 25 Hz in three age groups. Each circle represents the normalized group mean ± SEM. The number of cells and animals is as in **(C)**. There were no statistically significant differences between groups. **(F)** : As **(E)** but for trains of 20 stimuli at 100Hz. The number of cells and animals is as in **(D)**. There were no statistically significant differences between groups.

#### Changes of synaptic latency of phasic EPSCs during train stimulation

For the 25 Hz trains, in the P10 group a progressive increase of the synaptic latency occurred between the 1^st^ and the 20^th^ stimulus, with the latency being +0.63 ± 0.12 ms higher by the end than at the beginning of the train (Figure 10C). In P20 group, a brief decrease in the synaptic latency during the initial few pulses (by -0.14 ± 0.09 ms at the 5^th^ pulse) was followed by a mild increase (by 0.150 ± 0.081 ms at the end of the train) (Figure 10C). The difference between P10 and P20 animals was highly significant (p=1.28E-04 to p=0.045). In P50 animals, we observed a more pronounced change in the synaptic latency after the initial few pulses (by 0.205 ± 0.063 ms) than at the end of the train (by 0.029 ± 0.084 ms) (Figure 10C). However, while the difference between P50 and P10 animals was statistically significant almost throughout the whole train (p=2.36E-06 to p=0.017), P20 and P50 groups did not test as different at any point (p=0.095 to p=0.985).

For the 100 Hz trains, in P10 group a progressive increase of the synaptic latency of the phasic EPSC occurred during the first 10 pulses, with a +1.409 ± 0.042 ms increase at the maximum when compared with the beginning of the train (Figure 10D). Subsequently, the changes in the synaptic latency became more erratic as stimulation triggered fewer EPSCs. In the P20 animals, changes of the synaptic latency resembled those during the 25 Hz trains: a brief shortening of the latency (by -0.097 ± 0.074 ms) during the initial few pulses, was followed by a mild increase during the subsequent stimuli within the train (Figure 10D). In the P50 group, a shortening of the latency during the initial phase of the train (by -0.165 ± 0.067 ms at the 6^th^ pulse) was followed by a mild progressive increase during the later phase of the trains reaching an increase of +0.236 ± 0.103 ms at the maximum (Figure 10D). Changes of the synaptic latency in the P10 group were significantly different from the P20 and P50 groups throughout the entire train (p=5.12E-07 to p=0.042, P10 vs P20; p=2.60E-10 to p=0.030, P10 vs P50), while the differences between P20 and P50 groups were significant only at some stimuli.

The variance of the synaptic latency of EPSCs at neuron-OPC was increasing during the time-course of both 25 Hz and 100 Hz trains in all three age-groups of animals (Figure 10E-F), and the increases were comparable between the experimental groups. The few statistically significant differences appeared largely coincidental.

Taken together, train stimulation affects synaptic latency of the phasic EPSC at neuron-OPC synapses, and this effect is more pronounced in younger than in mature animals.

#### Heterogeneity in synaptic latency changes

We also sought to determine how uniform the changes in the synaptic latency were within our recorded OPC population. We divided the OPCs into five categories, based on the proportion of synaptic latency change during the 25 Hz stimulation, ranging from OPCs showing only increase to OPCs showing continuous decrease (see Materials and Methods) (Figure 11 A), and compared the ratios between groups.

**Figure 11:**
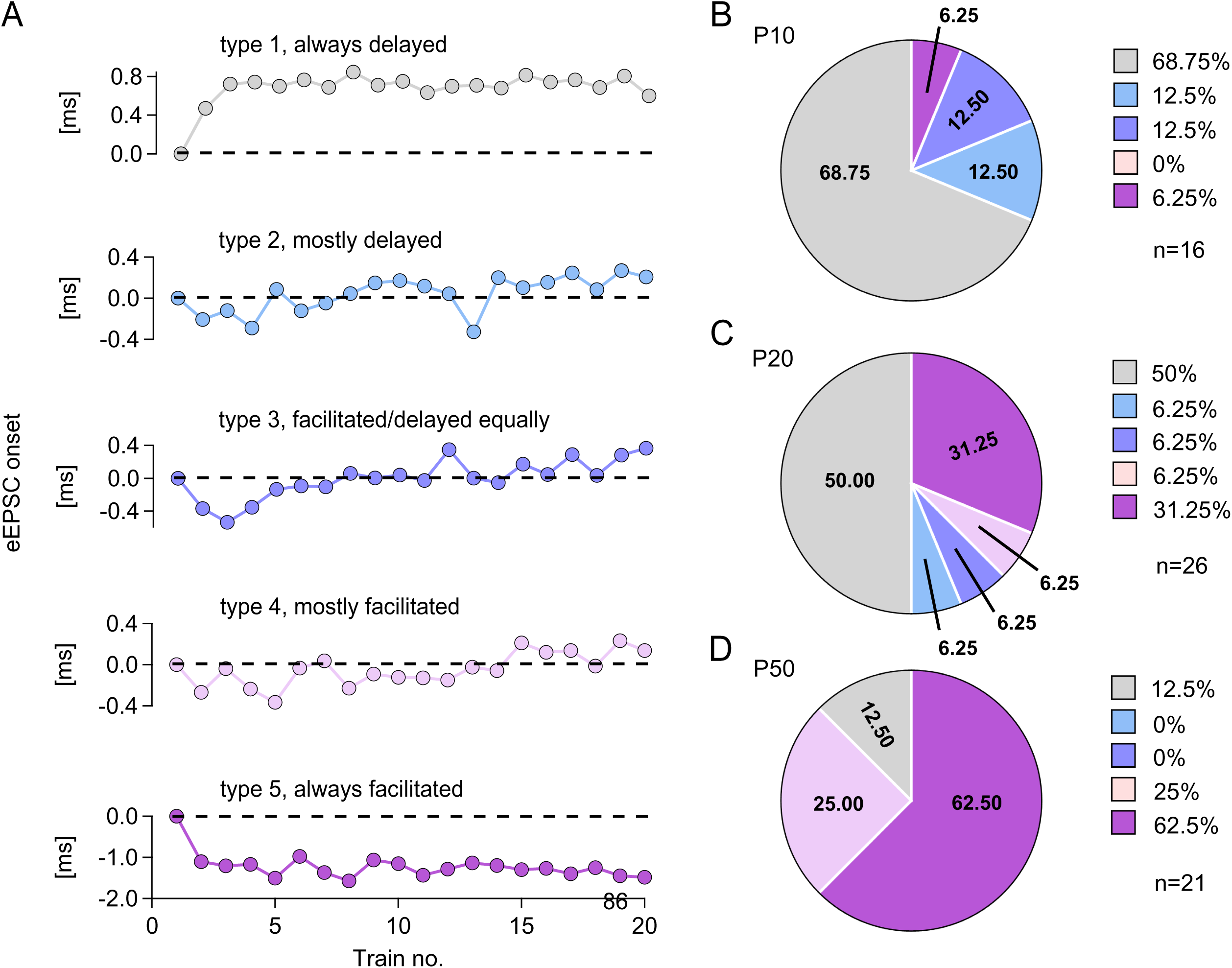
Heterogeneity of the synaptic latencies of phasic EPSCs at neuron-OPC synapses in the corpus callosum. **(A)** : Examples of five distinct categories based on the extent of synaptic latency change during 25 Hz stimulation, ranging from synapses showing only increase in latency to synapses showing continuous decrease in latency. Each example shows the normalized EPSC onset latency during the phasic (first 5 ms) component of the response after each stimulus in the train. **(B)** : Pie chart showing the percentage of OPCs in each of the five categories in the P10 group, color-coded to match the examples in (A). n = 16 cells from 14 animals. **(C)** : Same as in (B) for the P20 group (n = 26 cells from 21 animals). **(D)** : Same as in (B) for the P50 group (n = 21 cells from 18 animals).

In the P10 group, the majority of OPCs (68.75%) showed continuous increase in the synaptic latency during the time-course of the train, while the remaining OPCs showed either no change (12.5%), a small increase followed by a decrease (12.5%), or an increase (6.5%), (Figure 11B).

In the P20 group, 50% of OPCs showed continuous increase in the synaptic latency during the time-course of the train, 31.25% showed a decrease, while the remaining 18.75% of OPCs belonged to the categories between those two main types (Figure 11C).

In the P50 group, the vast majority of OPCs (62.5%) showed continuous decrease of the synaptic latency during the time-course of the train, 25% showed no/little change, while 12.5% of OPCs showed an increase (Figure 11D).

The P10 and P20 groups did not test as significantly different (p=0.070), but both tested as significantly different from P50 (P10 vs P50, p=3.95E-05; P20 vs P50, p=0.034; Fisher’s Exact test). Moreover, we found a strong and highly significant correlation between the synaptic latency category and the STP response shape during the 25 Hz trains in all cells, regardless of the age (r = 0.65, p = 1.93E-08), and in the P10 group (r = 0.670, p = 0.007; Spearman’s r). In the P20 group, the correlation was moderate but also significant (r = 0.472, p = 0.015; Spearman’s r). Surprisingly, in the P50 group the correlation was small and not significant (r = 0.143, p = 0.559; Spearman’s r), likely reflecting largely complete synapse maturation at this age.

### 2.9. Computational modelling suggests that both pre- and postsynaptic mechanisms may underlie STP at neuron-OPC synapses

To simulate potential mechanisms underlying the observed STP in OPCs from different age groups, we used the same OPC passive models established in Section 2.4. For each single synaptic event, we assumed that the total synaptic current is the sum of separate K^+^, Na^+^ and Ca^2+^ currents. The Ca^2+^ conductance depends on the age-dependent fraction, *f*_*ca*_, of Ca^2+^- permeable AMPARs. This parameter was set to match experimental findings (Figure 5): *f*_*ca*_ = 0.2 for P10; *f*_*ca*_ = 0.68 for P20; and *f*_*ca*_ = 0.74 for P50. Synaptic depression is modeled as a presynaptic mechanism. However, we consider two possible mechanisms underlying synaptic facilitation: a presynaptic mechanism and a postsynaptic mechanism.

#### Assumption 1: Solely presynaptic mechanisms underlie STP at neuron-OPC synapses

In the first set of simulations, we assumed that purely presynaptic mechanisms underlie STP. The primary synaptic parameters that differed between the age groups are *a*_*d*_ and *a*_*f*_, which are related to the rates of depression and facilitation, respectively (see Materials and Methods for details). In order to match with our experimental data, we set *a*_*d*_ and *a*_*f*_ as following: in the P10 group, *a*_*d*_ = .2 *and a*_*f*_ = 0 so there is no facilitation; in the P20 group, *a*_*d*_ = .03 and *a*_*f*_ = .33 so there is both depression and facilitation; and in the P50 group, *a*_*d*_ = 0 and *a*_*f*_ = .33 so there is no depression.

Note that the total synaptic conductance at the first input is *g*_*tot*_ = *g*_*K*_ + *g*_*Na*_ + *f*_*ca*_*g*_*ca*_. Since *f*_*ca*_ increases with age and *g*_*K*_, *g*_*Na*_ and *g*_*ca*_ are assumed to be age independent, it follows that *g*_*tot*_ increases with age. This is consistent with our experimental finding that the single-channel conductance of AMPARs in OPCs increases upon animal maturation.

For each simulation iteration, an excitatory synaptic connection was randomly placed along the OPC’s processes. When we set a model presynaptic neuron to fire at 100 Hz, the simulation result (Figure 12A bottom panels) closely recapitulated our experimental recordings of the somatic current (Figure 2) for the P10, P20 and P50 age groups. That is: in P10 animals, we observed short-term synaptic depression during the trains; in P20 animals, brief initial potentiation occurred during the first 4 stimuli of the train followed by depression; and in P50 animals, we observed strong synaptic potentiation with continuous increase in the amplitude reaching a plateau. We can then also simulate the somatic membrane potential (Figure 12A top panels). Figure 12B shows normalized somatic currents.

**Figure 12:**
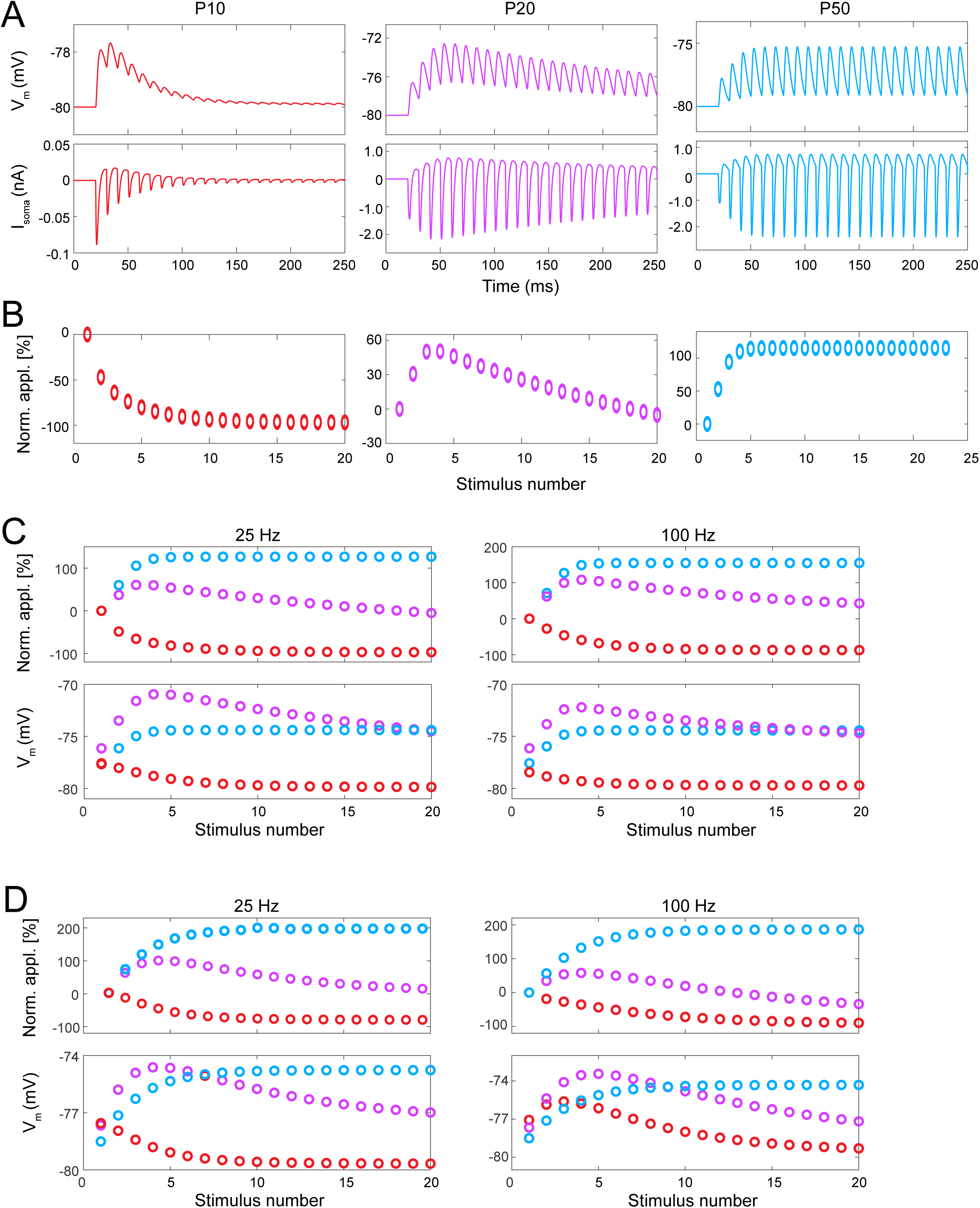
Computational modelling suggests that both pre- and postsynaptic mechanisms may underlie STP at neuron-OPC synapses. The same OPC passive models established in Figure 6 were used here. For each single synaptic event, instead of introducing a double-exponential *g*_*syn*_ with fixed rise and decay kinetics, we introduced *g*_*tot*_ as different ions’ conductance (Na^+^, K^+^ and Ca^2+^) through AMPARs: 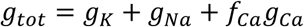 where *f*_*ca*_ represents the fraction of Ca^2+^-permeable AMPARs. **(A)** : We placed an excitatory synaptic connection randomly on each OPC model’s processes. We set the presynaptic neuronal firing rate at 100Hz. Here, we assume that STP we observed from experimental data is purely contributed by presynaptic mechanisms. Thus, for postsynaptic AMPAR’s conductance *g*_*tot*_, *g*_*K*_ and *g*_*Na*_ remain the same across age groups, where the Ca^2+^ conductance is determined by *f*_*ca*_*g*_*ca*_, and *f*_*ca*_undergoes age-dependent increase. By setting individual parameters controlling the magnitude of synaptic depression and facilitation (see Methods), our simulation of somatic synaptic current nicely recapitulates experimentally recorded STP across three age groups (bottom panel). In addition, we can simulate the somatic membrane potential during STP (top panel). **(B)** : Dot plots show normalized somatic currents. The normalization was executed as following: Let *M*_*k*_ be the maximum (in absolute value) somatic current in response to the k^th^ input. Then the normalized somatic currents are defined as (*M*_*k*_ − *M*_1_)⁄*M*_1_. **(C)** The simulations were repeated 25 times at two frequencies (25 Hz and 100 Hz) with synapses being placed at different randomly chosen locations. Dot plots demonstrate the average normalized somatic current amplitude (top panel) and average somatic membrane potential (bottom panel). **(D)** In a separate set of simulations, we assume that the STP is caused by a mixture of pre- and post-synaptic mechanisms. The presynaptic mechanism refers to presynaptic depression, and the postsynaptic facilitation takes place due to the unblocking of Ca^2+^-permeable AMPARs. The postsynaptic AMPAR conductance is determined by 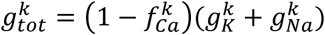 We ran 25 iterations of simulation using these parameters across three age groups at 25 Hz and 100 Hz. Similar as in **(C)**, dot plots demonstrate the average normalized somatic current amplitude (top panel) and average somatic membrane potential (bottom panel). Note that the simulation results for both **(C)** and **(D)** were very similar to our recorded experimental data.

We next considered average responses over multiple stimulations of multiple model OPCs within each group. The same twelve passive model OPC cells in each age group were used as in part 2.4. The simulations were repeated 25 times, with different randomly chosen synaptic locations for each model OPC, and for both 25 and 100 Hz presynaptic firing frequency. Finally, we took the average of the somatic current and membrane potential amplitudes over all simulations of all cells within each age group for each presynaptic firing frequency. We found that the modelled time-courses of synaptic current amplitude (Figure 12C) appear very similar to that recorded in the experimental setting (Figure 2), suggesting that the mechanism of STP at neuron-OPC synapses may indeed be presynaptic.

#### Assumption 2: Both pre- and postsynaptic mechanisms underlie STP at neuron-OPC synapses

In the second set of simulations, we assumed that the STP is caused by a mixture of pre- and post-synaptic mechanisms. The presynaptic mechanism refers to presynaptic depression and postsynaptic facilitation takes place due to the unblocking of Ca^2+^-permeable AMPARs. The parameters that change between age groups are the rate of depression, *a*_*d*_; the fraction of Ca^2+^- permeable AMPARs, *f*_*ca*_; and the maximal conductances of the K^+^ and Na^+^ currents, *g*_*K*_ and *g*_*Na*_. Values for *a*_*d*_ for the different age groups are given in Table 1 and values for *f*_*ca*_ were given previously.

Values of *g*_*K*_ and *g*_*Na*_ were chosen based on the experimental finding that single-channel conductance of AMPARs in OPCs increases upon animal maturation. For age k, where k = 10, 20 or 50, let 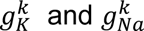 be the corresponding values of *g*_*K*_ and *g*_*Na*_, respectively. Then the total synaptic conductances for the 3 age groups at the first input is

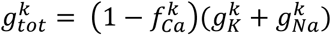

We assumed that the number of AMPAR channels remained constant across P10, P20 and P50 groups. Therefore, local synaptic conductance is proportional to single AMPAR channel conductance, which were determined experimentally. We have chosen each 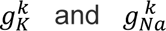 so that the ratios 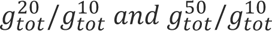 are the same as those for the corresponding single AMRAR conductance, i.e.1.1376 and 1.3474, respectively.

Using these simulation settings, we found that the modelled time-courses of synaptic current amplitudes (Figure 12D top panels) appeared very similar to that recorded in the experimental settings (Figure 2). The differences in STP response dynamics between the age groups was also reflected in the changes of the membrane potential responses (Figure 12D bottom panels). This suggests that the mechanism of STP at neuron-OPC synapses may also be postsynaptic.

Taken together, our computational modelling suggests that both pre- and postsynaptic mechanisms may contribute to the mechanisms of STP and to the switch from short-term depression to short-term facilitation during postnatal development, at neuron-OPC synapses in the corpus callosum.

## 3. Discussion

In this study, we have used experimental and computation approaches to investigate how maturation-dependent changes at neuron-OPC synapses in the mouse corpus callosum, putting a special focus on STP. Our major findings are: (1) During postnatal development of mice, there is a switch from short-term synaptic depression to short-term facilitation at neuron-OPC synapses; (2) Synaptic latency of phasic EPSC shortens, and the amount of asynchronous release decreases at neuron-OPC synapses as animals mature – those changes indicate that neurotransmitter release at neuron-OPC synapses becomes more synchronized during callosal maturation; (3) Postsynaptic AMPARs in callosal OPCs become more Ca^2+^-permeable as animals mature, and the changes are already observed at three weeks of the postnatal development; (4) Our computational modelling suggests that both pre- and postsynaptic changes may contribute to the developmental differences in the STP.

### 4.1. Functional significance of STP at neuron-OPC synapses

We found that train stimulation of callosal axons at 25 or 100 Hz triggers STP at neuron-OPCs synapses in the corpus callosum. Furthermore, STP switches from short-term synaptic depression to short-term synaptic facilitation during postnatal development. In behavioral paradigms *in vivo*, callosally-projecting neurons have been reported to fire at various frequencies (Caldwell et al., 2023; Ramos et al., 2008; Slater and Isaacson, 2020). Therefore, STP that we recorded at neuron-OPCs synapses is likely to occur under physiological conditions *in vivo* as well. What may be the functional significance of STP at neuron-OPC synapses, and its postnatal switch from depression to facilitation?

Several research groups, including ours, suggested that neuron-OPC synapses represent important communication sites through which neurons modulate development and function of the oligodendrocyte lineage cells, as well as myelination (Chen et al., 2018; Gibson et al., 2014; Kougioumtzidou et al., 2017; Kukley et al., 2008; Kukley et al., 2010; Nagy et al., 2017). Synaptic inputs are expected to trigger local and/or global cellular changes in the OPCs, including changes in the membrane potential, ion fluxes, etc. Our computational modelling suggests that, over the course of postnatal development, STP at neuron-OPC synapses contributes to a prolongation of the peak depolarization in OPCs. This prolonged depolarization may widen the temporal window for Ca^2+^-dependent signaling in OPCs and thereby facilitate the coupling of neuronal synaptic activity to various OPC behaviors. For example, AMPAR-mediated depolarization can trigger Ca^2+^ influx through voltage-gated Ca^2+^ channels, engaging Ca^2+^-dependent cascades such as CaMK and MAPK/ERK - pathways that influence proliferation and differentiation (Barron and Kim, 2019). These signals may also converge on PI3K-Akt and mTOR pathways, which are critical regulators of OPC survival and myelin production (Gaesser and Fyffe-Maricich, 2016; Suo et al., 2019; Waggener et al., 2013).

Thus, STP at neuron-OPC synapses may represent one of the phenomena which help neurons modulate behavior and/or functions of OPCs, in an activity-dependent way. The age-dependent switch from short-term depression to short-term potentiation, which we observed in the present study, may reflect developmentally regulated tuning of OPC sensitivity to neuronal activity. Short-term depression at synapses in the juvenile mice indicates that only few initial spikes in the train are “recognized” by OPCs and depolarize their membrane while the subsequent inputs are dampened. This may be important for preventing the response of OPCs to uncoordinated neuronal firing which likely still occurs in P10-P12 mice, because the refinement and pruning of callosal projections continues until the third postnatal week (Fame et al., 2011; Mizuno et al., 2010; Tagawa and Hirano, 2012). Switching to short-term potentiation and stronger membrane depolarization in more mature animals indicates that OPCs become more responsive to sustained repetitive firing of neurons. As facilitation enhances synaptic strength at neuron-OPC synapses, it favors the recognition of more active neurons by OPCs, e.g. those that fire with specific patterns and/or fire more frequently, and are more likely to be involved in experience-driven circuits. This may help OPCs to select more active axons and to prioritize their myelination, thus supporting the refinement of neuronal circuits.

Remarkably, many neuronal synapses also show age-dependent switch from short-term depression to short-term potentiation (Cheetham and Fox, 2010; Chen and Roper, 2004; Oswald and Reyes, 2008; Reyes and Sakmann, 1999). This switch usually reflects maturation of the presynaptic vesicular release machinery including changes in vesicular release probability, altered dynamics of synaptic vesicles, incorporation of all proteins important for exo- and endocytosis of synaptic vesicles, etc. (Andreae et al., 2012; Mozhayeva et al., 2002; Sudhof, 2012). A similar shift, from short-term depression to short-term potentiation, at neuron-OPC synapses suggests that OPCs are well integrated into the neuronal networks, and are able to adapt their processing of synaptic neuronal inputs to the maturation of the neuronal circuits. Furthermore, studying STP in the corpus callosum, we investigated synaptic release machinery in the axonal shafts of the same neurons whose synaptic boutons form neuronal synapses in the cortex. Our findings regarding changes in STP suggest that synaptic release machinery located at different sites along the same axon (i.e. axonal shaft in the white matter vs synaptic boutons in the grey matter) mature in a very similar fashion.

### 4.2. Is STP at neuron-OPC synapses of presynaptic or postsynaptic origin?

An important question usually asked by physiologists studying neurons is whether plasticity at synapses, be it STP or LTP, is of presynaptic or postsynaptic origin. A classical view for many synapses in the brain, has been that presynaptic mechanisms underlie STP, while postsynaptic mechanisms underlie LTP, but some postsynaptic mechanisms of STP and presynaptic mechanisms of LTP have been described as well, and many controversies still remain.

With regard to STP, at neuronal synapses the major presynaptic mechanisms mediating short-term synaptic facilitation are usually related to the prolonged increase in presynaptic Ca^2+^, which enhances the release probability of synaptic vesicles (Citri and Malenka, 2008; Fioravante and Regehr, 2011; Huganir and Nicoll, 2013; Regehr, 2012; Zucker and Regehr, 2002). This may be due to the accumulation of residual Ca^2+^, increase of presynaptic Ca^2+^ influx triggered by activity-dependent changes in the presynaptic Ca^2+^-channels, or saturation of the presynaptic endogenous Ca^2+^ buffers (e.g. calbindin) (Citri and Malenka, 2008; Fioravante and Regehr, 2011; Huganir and Nicoll, 2013; Regehr, 2012; Zucker and Regehr, 2002). The major presynaptic mechanisms mediating short-term synaptic depression include a progressive depletion of the synaptic vesicles pool available for release (Citri and Malenka, 2008; Fioravante and Regehr, 2011; Huganir and Nicoll, 2013; Regehr, 2012; Zucker and Regehr, 2002), and inactivation of presynaptic Ca^2+^ channels leading to reduced influx of Ca^2+^ required for triggering vesicle fusion and neurotransmitter release (Citri and Malenka, 2008; Fioravante and Regehr, 2011; Huganir and Nicoll, 2013; Regehr, 2012; Zucker and Regehr, 2002).

The major postsynaptic mechanisms contributing to short-term facilitation include polyamine-dependent facilitation of AMPARs (Rozov et al., 2018) and recruitment of silent AMPARs (Voronin and Cherubini, 2004), while major postsynaptic mechanisms contributing to short-term depression include AMPAR desensitization and receptor saturation, which may reduce synaptic response even when sufficient neurotransmitter is released (Chater and Goda, 2014).

Little is currently known about the mechanisms of STP at neuron-OPC synapses, and of its developmental switch from depression to facilitation. Our experimental findings show that in the juvenile mice, short-term synaptic depression during train stimulation goes along with a decrease in the response probability, an increase of the synaptic delay of phasic EPSCs, and an increase in the proportion of asynchronous release during the train. At neuronal synapses, a decrease in the response probability during the train is usually of presynaptic origin: it is attributed to the decline of the vesicular release probability due to the rapid depletion of the readily releasable pool (RRP) of synaptic vesicles, i.e. this pool is used up faster than it can be refilled (Fioravante and Regehr, 2011; Kaeser and Regehr, 2017). Synaptic latency at many neuronal synapses is inversely correlated with the release probability during train stimulation: depressing synapses tend to have increasing latencies, while facilitating synapses may show decreasing latencies (Boudkkazi et al., 2007). Similar mechanisms may operate at neuron-OPC synapses. Our results suggest that this indeed may be the case: if fewer vesicles are available in the RRP, neuron-OPC synapses may utilize vesicles from slower-releasing pools in order to support neurotransmitter release. This, in turn, may result in a larger synaptic delay of the phasic EPSC and occurrence of the asynchronous release, which we indeed observed.

Asynchronous release can be sustained for continued periods of time because it may utilize different presynaptic Ca^2+^ sources from those underlying the phasic release: while phasic release relies on the rapid precise transient influx of Ca^2+^ through voltage-gated Ca^2+^ channels at the active zone, asynchronous release is triggered by residual Ca^2+^ and/or Ca^2+^ from secondary sources e.g. intracellular Ca^2+^ stores or Ca^2+^ entry at the axonal locations other than active zones (Chamberland et al., 2020; Kaeser and Regehr, 2014; Wen et al., 2013). An increase in asynchronous release during the time-course of the train may be a mechanism that helps synapses to encode the firing pattern of incoming action potentials into a sustained increase in neurotransmitter release (Kaeser and Regehr, 2014). Interestingly, we observed that at neuron-OPC synapses the contribution of asynchronous release to the total neurotransmitter release was larger in the juvenile than in more mature animals. This may indicate that the release machinery in juvenile animals is immature, and some synaptic vesicles may require more extensive priming before docking leading to asynchronous release, while at the later developmental stages priming is faster, phasic release becomes more reliable, and asynchronous release decreases. At the same time, the presence of asynchronous release at synapses of juvenile animals and its increasing dynamics during the time-course of the train indicates that presynaptic vesicular release machinery in the juvenile animals is not compromised, in spite of the short-term depression of the phasic release. Phasic and asynchronous release may rely on two different pools of synaptic vesicles: the fast-releasing (which are located closer to the main Ca^2+^ sensor, are released fast, but are also depleted fast during the repetitive stimulation) and the slow-releasing vesicles (which are located more far away from the main Ca^2+^ sensor, and are released and depleted slower during the repetitive stimulation), respectively (Hagler and Goda, 2001; Schneggenburger et al., 2002; Tran et al., 2022; Trommershauser et al., 2003). Taken together, even though OPCs in juvenile mice might mainly “recognize” few initial spikes in the train, they may still be able to sense some amount of later release during repetitive stimulation.

Can postsynaptic mechanisms contribute to short-term depression in the juvenile mice as well? In our experiments, we have observed that depression of the phasic EPSC amplitude during the train was larger than the decrease in the response probability. Such a scenario may occur if postsynaptic AMPARs in OPCs get desensitized by an augmented amount of glutamate released during repetitive stimulation (Trussell et al., 1993). However, we think that this mechanism does not have a large impact in our case: if AMPARs were desensitized we would expect to see a decrease in the amplitude not only of phasic EPSCs, but also of asynchronous EPSCs, and a decrease in their frequency during the time-course of the trains. To the contrary, we have observed an increase in the amount of asynchronous release.

In P20 and P50 groups, we have observed short-term facilitation of phasic EPSCs during the trains, which occurred to various degrees and with various time-courses depending on the stimulation paradigm. The short-term facilitation was accompanied by an increase in the response probability during the train, and a decrease in the synaptic delay of the phasic EPSCs. Asynchronous release still occurred in the P20 and P50 groups, but it was less pronounced than in P10 group. Based on these findings, we think that during postnatal development glutamate release at neuron-OPC synapses gradually becomes more synchronized, and vesicular release machinery can better adapt to and sustain the repetitive neuronal activity.

Taken together, our experimental findings showing changes in the response probability, synaptic delay, and asynchronous release suggest that presynaptic mechanisms may contribute to STP at neuron-OPC synapses in the corpus callosum. Our computational modeling supports this idea showing that a switch from short-term depression to short-term facilitation can be indeed mediated by presynaptic mechanisms. This switch could be explained by maturation of both presynaptic voltage-gated calcium channels and RRPs, which have been suggested by reports studying neuronal synapses (Feldmeyer and Radnikow, 2009).

Interestingly, however, our computational modeling suggests that postsynaptic mechanisms may contribute to the switch from short-term depression to short-term facilitation, as well. When we considered an increased Ca^2+^ permeability of AMPARs and a greater morphological complexity of OPCs (Kula and Kukley, 2025) (also supported by an increase in C_m_) in the adult vs juvenile mice, the modelling results suggested that polyamine-dependent facilitation of AMPARs may be an additional mechanism contributing to the switch from short-term depression to short-term facilitation. During polyamine-dependent plasticity, activity-dependent changes in the synaptic strength are mediated by a progressive relief of the polyamine block on postsynaptic Ca^2+^-permeable AMPARs enhancing ion flux through the channels (Rozov et al., 2018). This facilitation during later developmental stages can counteract presynaptic depression or induce synaptic potentiation, acting as a postsynaptic mechanism. Although we have not specifically tested for this scenario in experimental settings, we consider it as an additional possible mechanism mediating a switch from short-term depression to short-term facilitation at neuron-OPCs synapses.

## 4. Conclusion

We demonstrated that during postnatal development of mice, there is a switch from short-term synaptic depression to short-term synaptic facilitation at neuron-OPC synapses in the corpus callosum, and neurotransmitter release at these synapses becomes more synchronized. Our computational modelling suggested that both pre- and postsynaptic changes may contribute to the developmental differences in STP. Intriguingly, several earlier studies found that similar paradigms of neuronal stimulation that induce plasticity at neuronal and neuron-OPC synapses can modulate various events involved in the myelination process, for instance, proliferation and differentiation of OPCs, and MBP synthesis, *in vitro* and *in vivo*. It is plausible that plasticity at neuron-OPC synapses underlies neuronal modulation of myelination, in analogy to as plasticity at neuronal synapses is thought to underlie learning and memory. Addressing this issue remains a challenge for future research.

## 5. Material and Methods

### 5.1. Ethics statement

All experiments were performed in accordance with current European Union guidelines and approved by the local government authorities for Animal Care and Use (Regierungspraesidium Tuebingen, State of Baden-Wuerttemberg, Germany). All efforts were made to minimize the suffering of the animals.

### 5.2. Animals

Heterozygous progeny of NG2DsRedBAC transgenic and C57BL/6 mice were used in this study. Breeding pairs of NG2DsRedBAC transgenic mice were originally obtained from the Jackson Laboratory (stock 008241). The C57BL/6 mice were originally obtained from Charles River. The mice were cross-bred in house under 12/12 hours light/dark conditions with water and food available *ad libitum*.

### 5.3. Slice preparation for electrophysiology

For patch-clamp recordings, we used coronal brain slices containing the central part of the corpus callosum, prepared from P8-11 (P10 group), P19-22 (P20 group) or P50-53 (P50 group) days-old mice of both sexes. Mice were anesthetized with a mixture of isoflurane in pure O_2_ (3% v/v) and decapitated. The brains were dissected in the ice-cold N-methyl-D-glucamine (NMDG)-based solution containing (in mM): 135 NMDG, 1 KCl, 1.2 KH_2_PO_4_, 20 choline bicarbonate, 10 glucose, 1.5 MgCl_2_, and 0.5 CaCl_2_ (pH 7.4, 310 mOsm), gassed with carbogen (95% O_2_, 5% CO_2_) for at least 30 min prior to the preparation. 300-μm-thick (for P7-10 animals), 270-μm-thick (for P19-22 animals) or 250-μm-thick (for P50-53 animals) coronal brain slices were cut in the same solution using Leica VT1200S vibratome. The slices were transferred to a Haas-type interface incubation chamber preheated to 32°C, and perfused with Ringer solution containing (in mM): 124 NaCl, 3 KCl, 1.25 NaH_2_PO_4_*H_2_O, 2 MgCl_2_, 2 CaCl_2_, 26 NaHCO_3_, 10 glucose; 300 mOsm/kg; 7.4 pH; gassed with carbogen. Afterwards, the slices were left to recover for ∼1 hour while the chamber was gradually cooling down to room temperature.

### 5.4. Patch-clamp recordings

After the recovery period, individual slices were transferred to the submerged recording chamber mounted on a stage of an upright microscope (FN-1, Nikon, Japan) equipped with infrared differential interference contrast (IR-DIC) filters and a fluorescence light source. Slices were superfused continuously (about 2 ml/min) with fresh carbogenated Ringer solution. All recordings were performed at room temperature (20-22°C). Data acquisition was performed in Igor Pro 6.3 (WaveMetrics, Lake Oswego, USA) using Recording Artist software written by Dr. Rick Gerkin (Arizona State University, USA).

#### 5.4.1. Identification of OPCs

OPCs were selected for recordings based on their strong red fluorescence, and were distinguished from pericytes based on their morphology and location away from the blood vessels.

Patch pipettes were pulled from borosilicate glass capillaries (Science Products, Germany) on a vertical puller (Model PC10, Narishige, Japan). Pipettes had resistance of 5-7 MOhms when filled with internal solution containing (in mM): 125 K-gluconate, 2 Na_2_ATP, 2 MgCl_2_, 0.5 EGTA, 10 HEPES, 20 KCl, 3 NaCl; 280–290 mOsm/kg, titrated to pH 7.3 with KOH. During all of the recordings, cells were voltage clamped at the holding potential V_h_ = −80 mV with an EPC-8 amplifier (HEKA, Germany). Liquid junction potential was calculated using the software JPCalc for Windows (Peter H. Barry, Sydney, Australia), and V_h_ was corrected by -13 mV before forming a seal (unless specified otherwise). Series resistance was not compensated. To verify that the selected cell was an OPC, after establishing the whole-cell configuration, 10 depolarizing voltage steps were applied from V_h_ = -80 mV in +10 mV increments. All recordings of currents evoked in response to voltage steps were low-pass filtered at 10 kHz and digitized with a sampling frequency of 20 kHz (ITC-18, HEKA Instruments Inc, USA). Cells without a clear OPC current profile were discarded.

#### 5.4.2. Recordings of the evoked synaptic currents

Evoked synaptic currents were elicited with the isolated pulse stimulator (A-M Systems, Model 2100, Science Products, Germany) using a glass pipette similar to the patch pipettes and filled with Ringer solution. The stimulation pipette was placed at the distance of 100 ± 25 μm from the recorded cell.

Biphasic rectangular pulses of 200–300 μs duration we applied. Single or paired (40 ms inter-pulse interval) were applied every 15s. Trains of stimuli (20 pulses at 25 or 100 Hz) were applied each 20 s.

To estimate release probability at neuron-OPC synapses, we used minimal stimulation paradigm in brain slices where the corpus callosum was isolated from the cortex by 4 cuts: 2 parallel and 2 perpendicular to the orientation of callosal axons (Nagy et al., 2017). We stimulated the callosal axons with voltage sufficiently low to activate one or at most few synaptic connections, and recorded phasic EPSCs in OPCs. In this paradigm, some stimulation trials yield an EPSC, while others do not trigger any response, allowing an estimation of the release probability at synapses. We performed 50-150 trials of stimulation, and analyzed the number of responses among all trials, i.e. the response probability = number of responses / number of trials.

All recordings were performed in the presence of NMDA-receptor antagonist (RS)-3-(2-Carboxypiperazin-4-yl)-propyl-1-phosphonic acid (CPP, 10 μM, Tocris) and GABA A-receptor antagonist SR95531 2-(3-Carboxypropyl)-3-amino-6-(4-methoxyphenyl) pyridazinium bromide (Gabazine, 5 μM, Sigma). To verify that synaptic currents were mediated by ionotropic glutamate receptors, we used 6-Cyano-7-nitroquinoxaline-2,3-dione (CNQX, 10 μM, Abcam), a blocker of all ionotropic glutamate receptors. All drugs were dissolved in Ringer solution and applied via the bath.

To monitor changes in the series resistance (R_s_) during recordings, we applied a 200-300 μs square test voltage step of -10 or -5 mV at the beginning of each sweep. All synaptic currents were low-pass filtered at 1 kHz and digitized with a sampling frequency of 10 kHz.

#### 5.4.3. Recordings of the I-V curves

For I-V curve recordings, we used a Cs-based internal solution containing (in mM): 100 CsCH_3_SO_3_H (CsMeS), 20 tetraethylammonium (TEA) chloride, 20 HEPES, 10 EGTA, 2 Na_2_ATP, and 0.2 NaGTP; 280-290 mOsm/kg; titrated to pH 7.3 with CsOH. As bath solution we used Ringer solution with elevated Ca^2+^ containing (in mM): 119 NaCl, 2.5 KCl, 1 NaH_2_PO_4_∗H_2_O, 1.3 MgCl_2_, 2.5 CaCl_2_, 26.2 NaHCO_3_, 11 glucose; 300 mOsm/kg; 7.4 pH; gassed with carbogen. Spermine, a potent voltage-dependent blocker of Ca^2+^-permeable AMPARs (Sigma, 100 μM) was included into the internal solution in all recordings of evoked EPSCs in order to test the Ca^2+^-permeability of AMPA receptors in OPCs. V_h_ was corrected for a −7 mV liquid junction potential before seal formation. The cells were held at different potentials (−90, −40, 0, +20, and +40 mV) and 10-50 sweeps were recorded at each potential. Monitoring of R_s_, and data filtering was done as described in 4.4.2.

#### 5.4.4. Criteria for discarding the recordings

All recordings included in this work had to meet three criteria in order to be included into the study: (1) The offset drift at the end of the experiment could not exceed ± 5 mV; and (2) R_s_ could not exceed 40 MΩ (all recorded cells had R_s_ between 20 and 40 MΩ); and (3) The change in R_s_ could not exceed ± 30% compared with the beginning of the experiment. The recordings which did not meet those criteria were discarded.

### 5.5. Analysis of the intrinsic properties of OPCs

Membrane resistance, R_m_, was calculated as the difference between the total resistance (input resistance, R_in_), obtained from the steady-state current in response to a small voltage step, and the series resistance, R_s_, obtained from the peak of the capacitive transient of the same step: R_m_ = R_in_ - R_s_.

Cell capacitance, C_m_, was estimated from the charge transferred into cell capacitance (ΔQ) by the following procedure. Let the change in the total current (ΔI(t)) denote the measured, time-dependent change in the current during application of ΔV. The capacitive current I_C_(t) was then calculated as the difference between the total and the resistive current (onset of voltage step at t = 0) as follows: the area under I_C_(t) approximates the change in charge transferred (ΔQ) and was transformed into cell capacitance, C_m_, with resting membrane potential (RMP,V_m_) = zero current potential.

The amplitude of the leak current (I_leak_) during a +10 mV step from the V_h_ = -80 (I_leak_ (+70mV)) was measured as the current amplitude at the end of the voltage step, i.e. when the current response was at the steady-state level. I_leak_ during the subsequent more positive voltage steps was calculated by multiplying I_leak_(+70mV) by a corresponding step factor, e.g. I_leak_(+60mV) = I_leak_ (+70mV) x 2; I_leak_ (+50mV) = I_leak_ (+70mV) x 3, etc.

Amplitude measurements of the voltage activated Na^+^ current (Na_v_), voltage activated fast A-type K^+^ current (K_v_ A-type), and voltage activated slow K^+^ current (K_v_ steady-state) were performed on the leak-subtracted current traces. The amplitude of the Na_v_ and K_v_ A-type was measured as the difference between the current peak and the baseline; the amplitude of the K_v_ steady-state was measured as the amplitude at the steady-state level and the baseline.

For each current type and for each cell, the current density was calculated by dividing the current amplitude by the C_m_ of the cell.

### 5.6. Analysis of the quantal EPSCs (qEPSCs)

While recording evoked EPSCs in OPCs elicited by train stimulation of callosal axons with 20 pulses at 25 or 100 Hz, we observed small delayed synaptic currents in OPCs occurring after cessation of stimulation. Based on the literature, and our previous study (Nagy et al., 2017), we considered those currents as quantal EPSCs.

For the analysis, we used quantal EPSCs with an onset of >10 ms after the last stimulus of the train. In each recorded OPCs, we collected the delayed EPSCs in 20-160 sweeps of 1.73-2.3 s length each. The EPSCs were detected using a deconvolution-based algorithm (Nagy et al., 2017; Pernia-Andrade et al., 2012) in FBrain, a customized program running under IgorPro 6 (WaveMetrics, Lake Oswego, USA). FBrain was kindly provided by Peter Jonas Lab (IST, Klosterneuburg, Austria). Additional digital high-pass (10 Hz) and Notch (50 ± 0.5 Hz) filtering was applied to the recorded sweeps in FBrain before the analysis. The convolved trace was passed through a digital band-pass filter at 0.001 to 200 Hz. The Pass 1 synthetic event detection template was constructed with a rise-time constant τ_rise_ = 0.5 ms, a decay time constant τ_decay_ = 4 ms, and an amplitude = −3 pA. The event detection threshold (θ) was set to 4.2 times the standard deviation of a Gaussian function fitted to the all-point histogram of the convolved trace, with with Freedman-Diaconis binning (Nagy et al., 2017; Pernia-Andrade et al., 2012). All events detected by the algorithm were inspected visually, and those events which clearly did not show kinetics of typical excitatory postsynaptic currents (i.e., fast rise time and exponential decay) were manually removed. All cells with removal ration > 30% were rejected from further analysis. The subsequent analysis was performed using custom-written macros in IgorPro.

### 5.7. Non-stationary fluctuation analysis (NSFA)

To estimate the single channel conductance of synaptic AMPARs in OPCs we performed peak-scaled non-stationary fluctuation analysis, NSFA (Hartveit and Veruki, 2007) as we have done previously (Chen et al., 2018; Nagy et al., 2017). In each cell, we inspected the delayed EPSCs visually, and selected for NSFA only the events with smooth rise- or decay phase. We applied the Spearman’s rank-order correlation test to validate that there was no change in the peak amplitude, rise-time, or decay-time of the EPSCs during the time-course of each experiment. We also made sure that there was no correlation between rise-time and decay-time constant (Hartveit and Veruki, 2007). If any of those correlations was found, we did not include the cell into the NSFA.

In the P10 group, we selected 13 EPSCs randomly from 8 cells; in the P20 and P50 groups, we selected 26 EPSCs randomly from 4 cells in each group. We pooled the selected EPSCs within each age group, resulting in 104 EPSCs per group included into the NSFA.

We aligned the selected EPSCs on the point of their steepest rise, and calculated the mean waveform from them. Subsequently, we scaled the amplitude of the mean waveform to the amplitude of each individual event. We then subtracted the scaled mean waveform from each individual event. This procedure generated the noise component for each event, from which we calculated the variance. The background variance, estimated from the segment of the trace before the onset of each event, was subtracted. The ensemble background-subtracted variance was calculated as an average of variances of all events.

We binned the mean amplitude wave into 8-10 bins and, based on this binning, we subsequently obtained the corresponding values of the binned variance (Hartveit and Veruki, 2007). We also tested whether varying the numbers of bins affected the results, but this was not the case. Finally, we plotted the values of the mean current variance versus the corresponding values of the mean current amplitude. The resulting variance-mean relationship was fitted with the following parabola function in IgorPro:

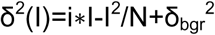

where *δ(I)* is the variance; *i* is the (weighted) estimate of a mean single channel current; *N* is average number of channels opened at the peak, and *δ_bgr_*is the background variance.

The single-channel conductance (γ) of synaptic AMPARs was calculated from the single-channel current i as:

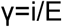

where E is the driving force for AMPAR-mediated EPSC, which is -80 mV in our study.

Notably, we cannot get the information about the total number of AMPAR channels from our NSFA analysis because we used the peak-scaling method for NSFA where the information about the number of ion channels is lost (Hartveit and Veruki, 2007).

All routines for NSFA were custom programmed and were based on the code presented in the supplementary material of (Hartveit and Veruki, 2007).

### 5.8. Analysis of the EPSCs evoked by single and paired stimulation, and the I-V curve

All analysis has been performed on the sweeps with subtracted stimulus artifacts. To remove the stimulus artifacts, we subtracted an average sweep containing no response from each EPSC response. To generate an average sweep containing no response to a single stimulus, we averaged sweeps containing EPSC failures after the first stimulus, or sweeps recorded in the presence of tetrodotoxin citrate or CNQX, or sweeps recorded at the holding potential of V_h_ = 0. To generate an average sweep containing no response to a paired stimulation, the averaged sweep with no response was concatenated with itself.

To measure the EPSC amplitude after first or second pulse, we used the following procedure: For each recorded sweep the baseline was adjusted to the 500 ms segment immediately preceding the stimulation; the peak-center of each event was determined as the time-point at which the first derivative of the sweep crossed zero; the amplitude values of the current at the peak ± 2 points around it were averaged, and the resulting value was taken as EPSC amplitude. The threshold for event detection was determined individually for each recorded sweep, and was equal to three times the standard deviation of the noise. In case several EPSCs occurred after a given stimulus, care was taken to measure the amplitude of the first event.

Although we used paired-pulse stimulation in all experiments, in order to generate the I-V curve we considered only the EPSCs occurring after the first pulse. The amplitudes of all recorded sweeps (10 to 15) at a given holding potential (−90, −40, 0, +20, +40 mV, and back to −90 mV) were measured as described above, and averaged. The resulting averages were used to generate the I-V curve in each cell.

To calculate the rectification index (RI), the average value of the EPSC amplitude at +40 mV was divided by the average value of the EPSC amplitude at −90 mV. RI is a measure of linearity of the I-V curve. In our experimental conditions, RI is expected to be 0.45 for a completely linear I-V curve (reflecting the presence of edited GluA2 subunit within AMPAR complex), while a decrease in the RI value describes inward rectification (reflecting the decrease or absence of the edited GluA2)(Coombs and Cull-Candy, 2021).

To determine the paired-pulse ratio, the average amplitude value of the EPSC occurring after the second pulse was divided by the average amplitude value of the EPSC occurring after the first pulse at a holding potential of −90 mV.

All analysis of the evoked EPSCs was performed using custom-written macros in IgorPro.

### 5.9. Estimation of the fraction of rectifying AMPARs (FRR) based on rectification measurements

To obtain the FFR, we used the equation developed by Stubblefield and Benke (Stubblefield and Benke, 2010), which allows for estimation of the FRR based on rectification measurements. Their rectification indices were calculated as a ratio of EPSC amplitudes recorded at −70 mV and at +40 mV. To match their approach, we took the inverse of our RI values (1/RI) from Figure 2E and adjusted their equation to model our results as follows:

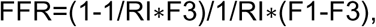

where F1 is the maximal block of inwardly rectifying receptors (EPSC at +40 mV/ EPSC at −90mV), extrapolated to be 0.035 (Chen et al., 2018). F3 is the value for linear relationship (F3 = 40/90 = 0.444). This analysis assumes that there is no change in presynaptic function.

### 5.10. Analysis of the EPSCs evoked by train stimulation

All analysis has been performed on the sweeps with subtracted stimulus artifacts. Stimulus artifact was subtracted using the following procedure: for each recorded cell, the sweeps containing failure in response to the first stimulus in the train were collected and averaged (failure was defined as an absence of postsynaptic response after a stimulation pulse); the part of the averaged sweep from the beginning of the first stimulus in the train until the end of the first inter-stimulus interval was cut out; a new sweep (“train of failures”) was generated by concatenating the cut piece of the train *n* times, in which *n* = number of stimuli in the train; the “train of failures” was subtracted from each recorded sweep.

The events were detected using a deconvolution-based algorithm in FBrain (Nagy et al., 2017; Pernia-Andrade et al., 2012) in the stimulus-artifact-subtracted sweeps containing a 1-s-long pre-train baseline.

We defined phasic EPSCs during the train as those events for which the EPSC onset was located ≤ 6 ms after the stimulus onset. We defined asynchronous EPSCs during the train as those events for which the EPSC onset was located within > 5 ms after the stimulus onset.

To estimate the amplitude of each phasic EPSC, we used the following procedure: the baseline was adjusted to the time interval from the beginning of each stimulus until the EPSC onset; the peak center of each EPSC was determined as a minimum (i.e., where the first derivative of the sweep crosses zero) within the interval of 7.5 ms after the stimulus onset; the values of the current in the peak center and in 2 points around it were averaged in order to obtain the measurement of the current amplitude. Hence, response was defined in our study as an EPSC for which onset is located ≤ 6 ms after the stimulus and whose peak is located ≤ 7.5 ms after the stimulus.

To estimate the amplitude of each failure, a similar procedure was used, with the exception that “the peak center” was determined not as a minimum but as one point randomly selected by the algorithm within the interval of 7.5 ms after the stimulus onset. Hence, a failure was defined in our study as a situation in which, after a stimulation pulse, we did not observe an EPSC with an onset located ≤ 6 ms after the stimulus and with peak located ≤ 7.5 ms after the stimulus.

For each stimulation paradigm in each cell, the mean amplitude of the phasic EPSCs after a given stimulus was calculated as the mean amplitude of all responses and all failures after this stimulus. Response probability was calculated as the number of responses after a stimulus divided by the total number of trials in the stimulation paradigm. To calculate the average amplitude and the response probability (after a given stimulus) across all cells for a given stimulation paradigm, the corresponding mean values were averaged. To calculate the normalized average amplitude of the EPSCs and the response probability, the corresponding average value after a given stimulus was divided by the average value after the first stimulus. To study the kinetics of phasic EPSCs, 20%–80% rise time and weighted decay time constant (after double-exponential fit) were measured for each event.

The latency of each phasic EPSC was determined as the time difference between the response onset (point where the convolved trace crossed the event detection threshold in FBrain) and the first time point of the stimulus. The baseline was adjusted to the time interval from the beginning of each stimulus until the EPSC onset; hence, only the transient currents after each stimulus were taken into consideration. To study changes in the synaptic latency, we analyzed the change in the time-interval between the start of the axonal stimulation and the normalized onset of the phasic EPSC after each pulse during the trains of 20 pulses at 25 or 100 Hz. For normalization, we subtracted the time-interval between the beginning of the stimulus and the median EPSC onset at the first stimulus in the train from all other measurements.

To estimate the charge transferred during phasic EPSCs, we split each inter-stimulus interval into 5-ms-long bins and performed trapezoidal integration on each bin. To calculate the total charge transfer during the train, the charge values after each stimulus within the train were summed.

### 5.11. Statistics

All data acquisition was randomized. Throughout the study we made all efforts to avoid pseudo-replication by restricting the maximum number of cells acquired from a single animal to 3. The exact number of cells and animals used in each experiment is given in the figure legends.

Statistical analysis was performed in Graph Pad Prism 9.3.1. All datasets were tested for homoscedasticity and normality. If the datasets had both normal distributions and equal variances, one-way ANOVA with post hoc Holm-Sidak’s test was used. If the datasets had normal distributions but unequal variances, Brown-Forsythe and Welch ANOVA with post hoc Dunnet’s T3 test were used. If the datasets were not normally distributed, but had equal variances, Kruskal-Wallis test with post hoc Dunn’s test were used. If the datasets were not normally distributed and had unequal variances, Brown-Forsythe and Welch ANOVA with post hoc Dunnet’s T3 test were used. For all statistical comparisons, significance level was set at p < 0.05. Statistically significant differences are indicated on the figures by star markings: ∗ represents p ≤ 0.05, ∗∗ represents p ≤ 0.01, and ∗∗∗ represents p ≤ 0.001. The exact p values are given in the text. If the data is normally distributed the data in the text or figures is reported as scatter plots and mean ± standard error of the mean (SEM) or, if not normally distributed as a boxplot containing median and 10th, 25th, 75th, 90th or 25th, 75th percentiles.

The datasets were checked for outliers by Prism’s ROUT method at Q=5%.

### 5.12. Power and sample size calculations

The sample sizes for all of the statistical comparisons in this work were determined based on the means and pooled standard deviations from preliminary recordings of 8-10 cells or slices per group, *α* = 0.05, *β* = 0.8, corrected for the number of pairwise comparisons (*τ*), based on the following equations:

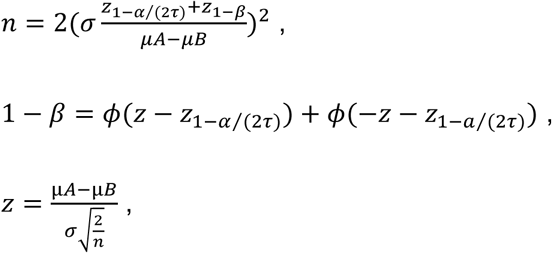

Where *n* is the sample size; *σ* is standard deviation; *ϕ* is standard Normal distribution function; *α* is Type I error; *τ* is the number of pairwise comparisons; *β* is Type II error. During the calculations, the normality of residuals and equality of variances were assumed *a priori*.

The calculations were performed in an online calculator available at: http://powerandsamplesize.com/Calculators/Compare-k-Means/1-Way-ANOVA-Pairwise

### 5.13. Computational modelling

#### 5.13.1. Passive cell model

To consider the complex highly ramified morphology of OPC, in our modelling we used the morphological parameters of those OPCs which we have analyzed in detail and described previously (Kula and Kukley, 2025). The reconstructed morphology data was imported into MATLAB to construct the network topology and then incorporated into the Neuron Simulation Environment for creating passive cell models. Twelve passive OPC models in each age group were created. The diameter of all OPC processes was set to be 0.2 µm and the diameter for the OPC soma to be 6.5 µm (Sun and Dietrich, 2013). The membrane properties in our passive models were set the same as described previously with specific membrane resistance, R_m_= 8000 Ω cm^2^, specific membrane capacitance, c_m_ = 1 µF/ cm^2^, and specific axial resistance, R_a_ = 100 Ω/cm (Sun and Dietrich, 2013).

We set the leak conductance to be consistent with our experimental data that the OPC input resistance (R_INP_) decreases and gradually becomes leakier, and cell surface area (SA) increases as mice grow older. Previously, Sun and Dietrich (Sun and Dietrich, 2013) used the specific membrane resistance R_m_ = 8000 Ω cm^2^ in a “ball-and-stick” model; this translates to the leak conductance, g_L_ = .000125 S cm^-2^. Here we use the same g_L_ = .000125 S cm^-2^ for the P10 group, and calculated the leak conductance for the P20 and P50 groups as follows. For K = 10, 20 or 50, let 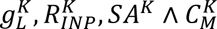 denote the average leak conductance, input resistance, surface area and total capacitance of the cells within age group PK. Then, for each K, 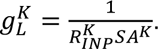 It follows that

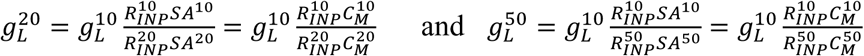

From the experimental data: 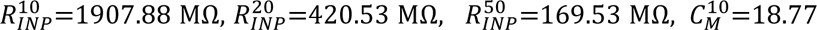 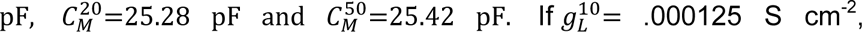 then from the above formulas: 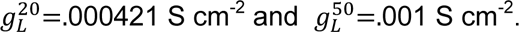

#### 5.13.2. Synaptic connection

We consider two separate models for the synaptic current. When considering the OPC response to a single synaptic input, we used a model for the synaptic current of the form *I*_*syn*_ = *g*_*syn*_*s*(*t*)(*V* − *E*_*syn*_). The model was implemented using NetStim within NEURON with *E*_*syn*_ = 0 mV and *s*(*t*) a double exponential with rise time *τ*_1_ = .25 ms and decay time *τ*_2_ = 1 ms.

A more detailed synaptic current model is needed when we study STP. We then assume that there is a model neuron that fires periodically and makes a glutamatergic synaptic connection within the OPC model network. We consider both Ca^2+^-permeable and non-Ca^2+^-permeable AMPARs. Each non-Ca^2+^-permeable receptor is permeable to K^+^ and Na^+^, while each Ca^2+^-permeable receptor is permeable to K^+^, Na^+^ and Ca^2+^. For convenience, we assume that the maximum conductance of each K^+^ current and each Na^+^ current through each AMPAR, whether it is Ca^2+^-permeable or non-Ca^2+^-permeable, is the same.

The model includes both short-term synaptic depression and synaptic facilitation. Synaptic depression is modeled as a presynaptic mechanism. However, we explore two possible mechanisms underlying synaptic facilitation: a presynaptic mechanism and a postsynaptic mechanism.

*Presynaptic mechanism:* Denote the fraction of Ca^2+^-permeable AMPARs as*f*_*ca*_. Based on our experimental findings showing age-dependent changes in the Ca^2+^-permeability of AMPARs in OPCs (Figure 5), we assume that the parameter *f*_*ca*_ changes with the age of the animals. The total synaptic current can then be written as

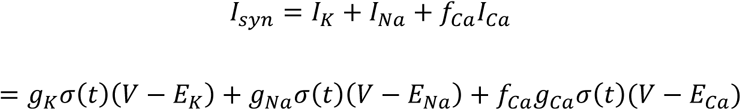

Here, *g*_*K*_ ∧ *g*_*Na*_ are maximum K^+^ and Na^+^ conductance, *g*_*ca*_ is the maximum Ca^2+^ conductance if all AMPARs are Ca^2+^-permeable, and *E*_*K*_, *E*_*Na*_ *and E*_*ca*_ are reversal potentials. We assume that

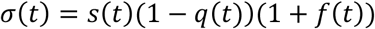

where *s*(*t*) is the fraction of open channels, and *q*(*t*) ∧ *f*(*t*) correspond to the magnitude of synaptic depression and facilitation, respectively. Assume that 0 ≤ *q*(*t*) ≤ 1 and *q*(0) = 0. If *q* = 0, then there is no depression, while if *q* = 1, then there is maximum depression. Furthermore, 0 ≤ *f*(*t*) ≤ *f*_*max*_ ∧ *f*(0) = 0. If *f* = 0, then there is no facilitation, while if *f* = *f*_*max*_ then there is a maximum amount of facilitation.

The basic idea is that when the presynaptic neuron fires a spike, *s*(*t*) approaches 1, and both *q*(*t*) and *f*(*t*) increase by certain amounts. Between spikes, each of these variables relaxes back to 0, at specified time rates. More precisely, these variables satisfy first order equations of the form

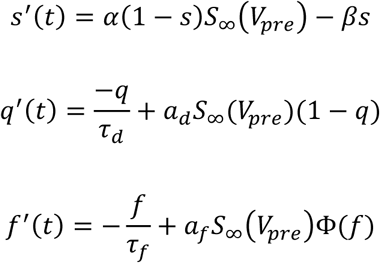

where *V*_*pre*_ is the presynaptic neuron’s membrane potential,

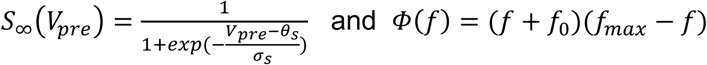

The function 𝛷(*f*) was chosen to match experimental data. Note that if *a*_*d*_ = 0 *or a*_*f*_ = 0, then there is no depression or facilitation, respectively. Based on the time-courses of STP in our experimental data (Figure 2), in the computer simulations we assumed that the parameters *a*_*d*_ *and a*_*f*_ change with animal age. Parameter values are listed in Table 1.

*Postsynaptic Mechanism:* As before, denote the fraction of Ca^2+^-permeable AMPARs as *f*_*ca*_, which we assume is age dependent. Moreover, we now assume that the Ca^2+^-permeable AMPARs may be blocked or unblocked, e.g., by endogenous polyamines. The unblocking of Ca^2+^-permeable AMPARs is assumed to be activity dependent, and this is what underlies postsynaptic facilitation. We also assume that the maximal conductances of the K^+^ and Na^+^ currents are age dependent. As before, there is age dependent presynaptic depression.

Let *γ*(*t*) equal to the fraction of Ca^2+^-permeable AMPARs that are unblocked. Then the fraction of all AMPARs that are unblocked is 1 − *f*_*ca*_(1 − *γ*). The total synaptic current can then be written as *I*_*syn*_ = *I*_*K*_ + *I*_*Na*_ + *I*_*ca*_, where

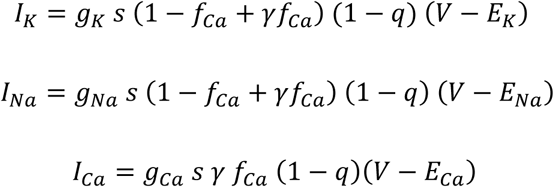

As before *q* corresponds to the magnitude of synaptic depression and *s* is the fraction of open channels. Both *q*(*t*) and *s*(*t*) satisfy the same equations as before.

The fraction, *γ*(*t*), of unblocked Ca^2+^-permeable AMPARs depends on the postsynaptic – that is, the OPC’s – membrane potential, which we denote as *V*_*post*_. We assume that *γ*(*t*) satisfies the equation

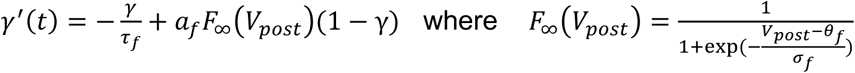

and *γ*(0) = 0. Note that *F*_∞_ is approximately the Heaviside step function. Whenever *V*_*post*_ increases past the threshold *θ*_*f*_, *γ* increases at the approximate rate *a*_*f*_(1 − *γ*). When *V*_*post*_ falls below *V*_*post*_, *γ* relaxes back towards 0 at the rate 1⁄*τ*_*f*_.

## Acknowledgments

We thank Rick Gerkin (Arizona State University, USA) for help with setting up the acquisition software, and Alejandro Pernía-Andrade (IST, Klosterneuburg, Austria) for help with setting up FBrain.

---

**Table:** STP Model Parameters.

|  |  |  |
| --- | --- | --- |
| $E_K = -90 \text{ mV}$ | $E_{Na} = 90 \text{ mV}$ | $E_{Ca} = 120 \text{ mV}$ |
| $\theta_s = -10 \text{ mV}$ | $\sigma_s = 5$ | $\tau_f = 500 \text{ ms}$ |
| $\tau_d = 1000 \text{ ms}$ | $\alpha = 4 \text{ ms}^{-1}$ | $\beta = .4 \text{ ms}^{-1}$ |

Age dependent parameters
| Presynaptic: P10; P20; P50 | Postsynaptic: P10; P20; P50 |
| --- | --- |
| $a_d$: .2; .03; 0 | $a_d$: .2; .05; 0 |
| $a_f$: 0; .33; .33 | $f_{Ca}$: .2; .68; .74 |
| $f_{Ca}$: .2; .68; .74 | $g_K = g_{Na}$: .000093 $\mu\text{S}$<br>.000264 $\mu\text{S}$<br>.000385 $\mu\text{S}$ |

Age independent parameters
| Presynaptic | Postsynaptic |
| --- | --- |
| $f_0 = .15$ | $a_f = .05$ |
| $f_{max} = 3$ | $\theta_f = -77 \text{ mV}$ |
| $g_K = .00018 \mu\text{S}$ | $\sigma_f = .5$ |
| $g_{Na} = .00018 \mu\text{S}$ | |
| $g_{Ca} = .00004 \mu\text{S}$ | |

## References

Andreae, L.C., Fredj, N.B., and Burrone, J. (2012). Independent vesicle pools underlie different modes of release during neuronal development. The Journal of neuroscience : the official journal of the Society for Neuroscience 32, 1867–1874.

Arguello, P.A., and Gogos, J.A. (2012). Genetic and cognitive windows into circuit mechanisms of psychiatric disease. Trends in neurosciences 35, 3–13.

Barron, T., and Kim, J.H. (2019). Neuronal input triggers Ca(2+) influx through AMPA receptors and voltage-gated Ca(2+) channels in oligodendrocytes. Glia 67, 1922–1932.

Boudkkazi, S., Carlier, E., Ankri, N., Caillard, O., Giraud, P., Fronzaroli-Molinieres, L., and Debanne, D. (2007). Release-dependent variations in synaptic latency: a putative code for short-and long-term synaptic dynamics. Neuron 56, 1048–1060.

Boudkkazi, S., Fronzaroli-Molinieres, L., and Debanne, D. (2011). Presynaptic action potential waveform determines cortical synaptic latency. The Journal of physiology 589, 1117–1131.

Caldwell, M., Ayo-Jibunoh, V., Mendoza, J.C., Brimblecombe, K.R., Reynolds, L.M., Zhu Jiang, X.Y., Alarcon, C., Fiore, E., J, N.T., Phillips, G.R., et al. (2023). Axo-glial interactions between midbrain dopamine neurons and oligodendrocyte lineage cells in the anterior corpus callosum. Brain structure & function 228, 1993–2006.

Chamberland, S., Timofeeva, Y., Evstratova, A., Norman, C.A., Volynski, K., and Toth, K. (2020). Slow-decaying presynaptic calcium dynamics gate long-lasting asynchronous release at the hippocampal mossy fiber to CA3 pyramidal cell synapse. Synapse 74, e22178.

Chater, T.E., and Goda, Y. (2014). The role of AMPA receptors in postsynaptic mechanisms of synaptic plasticity. Frontiers in cellular neuroscience 8, 401.

Cheetham, C.E., and Fox, K. (2010). Presynaptic development at L4 to l2/3 excitatory synapses follows different time courses in visual and somatosensory cortex. The Journal of neuroscience : the official journal of the Society for Neuroscience 30, 12566–12571.

Chen, H.X., and Roper, S.N. (2004). Tonic activity of metabotropic glutamate receptors is involved in developmental modification of short-term plasticity in the neocortex. Journal of neurophysiology 92, 838–844.

Chen, T.J., Kula, B., Nagy, B., Barzan, R., Gall, A., Ehrlich, I., and Kukley, M. (2018). In Vivo Regulation of Oligodendrocyte Precursor Cell Proliferation and Differentiation by the AMPA-Receptor Subunit GluA2. Cell reports 25, 852–861 e857.

Citri, A., and Malenka, R.C. (2008). Synaptic plasticity: multiple forms, functions, and mechanisms. Neuropsychopharmacology : official publication of the American College of Neuropsychopharmacology 33, 18–41.

Coombs, I.D., and Cull-Candy, S.G. (2021). Single-channel mechanisms underlying the function, diversity and plasticity of AMPA receptors. Neuropharmacology 198, 108781.

Crabtree, G.W., and Gogos, J.A. (2014). Synaptic plasticity, neural circuits, and the emerging role of altered short-term information processing in schizophrenia. Frontiers in synaptic neuroscience 6, 28.

Cull-Candy, S., Kelly, L., and Farrant, M. (2006). Regulation of Ca2+-permeable AMPA receptors: synaptic plasticity and beyond. Current opinion in neurobiology 16, 288–297.

De Leon Reyes, N.S., Bragg-Gonzalo, L., and Nieto, M. (2020). Development and plasticity of the corpus callosum. Development 147.

Del Castillo, J., and Katz, B. (1954). Quantal components of the end-plate potential. The Journal of physiology 124, 560–573.

Deng, P.Y., Sojka, D., and Klyachko, V.A. (2011). Abnormal presynaptic short-term plasticity and information processing in a mouse model of fragile X syndrome. The Journal of neuroscience : the official journal of the Society for Neuroscience 31, 10971–10982.

Enyedi, P., and Czirjak, G. (2010). Molecular background of leak K+ currents: two-pore domain potassium channels. Physiological reviews 90, 559–605.

Etherington, S.J., and Williams, S.R. (2011). Postnatal development of intrinsic and synaptic properties transforms signaling in the layer 5 excitatory neural network of the visual cortex. The Journal of neuroscience : the official journal of the Society for Neuroscience 31, 9526–9537.

Falk, T., Kilani, R.K., Strazdas, L.A., Borders, R.S., Steidl, J.V., Yool, A.J., and Sherman, S.J. (2003). Developmental regulation of the A-type potassium-channel current in hippocampal neurons: role of the Kvbeta 1.1 subunit. Neuroscience 120, 387–404.

Fame, R.M., MacDonald, J.L., and Macklis, J.D. (2011). Development, specification, and diversity of callosal projection neurons. Trends in neurosciences 34, 41–50.

Feldmeyer, D., and Radnikow, G. (2009). Developmental alterations in the functional properties of excitatory neocortical synapses. The Journal of physiology 587, 1889–1896.

Feng, L., Molnar, P., and Nadler, J.V. (2003). Short-term frequency-dependent plasticity at recurrent mossy fiber synapses of the epileptic brain. The Journal of neuroscience : the official journal of the Society for Neuroscience 23, 5381–5390.

Fields, R.D. (2008). White matter in learning, cognition and psychiatric disorders. Trends in neurosciences 31, 361–370.

Fioravante, D., and Regehr, W.G. (2011). Short-term forms of presynaptic plasticity. Current opinion in neurobiology 21, 269–274.

Gaesser, J.M., and Fyffe-Maricich, S.L. (2016). Intracellular signaling pathway regulation of myelination and remyelination in the CNS. Experimental neurology 283, 501–511.

Gibson, E.M., Purger, D., Mount, C.W., Goldstein, A.K., Lin, G.L., Wood, L.S., Inema, I., Miller, S.E., Bieri, G., Zuchero, J.B., et al. (2014). Neuronal activity promotes oligodendrogenesis and adaptive myelination in the mammalian brain. Science 344, 1252304.

Giovedi, S., Corradi, A., Fassio, A., and Benfenati, F. (2014). Involvement of synaptic genes in the pathogenesis of autism spectrum disorders: the case of synapsins. Frontiers in pediatrics 2, 94.

Goldstein, S.A., Bockenhauer, D., O’Kelly, I., and Zilberberg, N. (2001). Potassium leak channels and the KCNK family of two-P-domain subunits. Nature reviews Neuroscience 2, 175–184.

Greger, I.H., Watson, J.F., and Cull-Candy, S.G. (2017). Structural and Functional Architecture of AMPA-Type Glutamate Receptors and Their Auxiliary Proteins. Neuron 94, 713–730.

Hagler, D.J., Jr., and Goda, Y. (2001). Properties of synchronous and asynchronous release during pulse train depression in cultured hippocampal neurons. Journal of neurophysiology 85, 2324–2334.

Haroutunian, V., Katsel, P., Roussos, P., Davis, K.L., Altshuler, L.L., and Bartzokis, G. (2014). Myelination, oligodendrocytes, and serious mental illness. Glia 62, 1856–1877.

Hartveit, E., and Veruki, M.L. (2007). Studying properties of neurotransmitter receptors by non-stationary noise analysis of spontaneous postsynaptic currents and agonist-evoked responses in outside-out patches. Nature protocols 2, 434–448.

Huganir, R.L., and Nicoll, R.A. (2013). AMPARs and synaptic plasticity: the last 25 years. Neuron 80, 704–717.

Jerng, H.H., Pfaffinger, P.J., and Covarrubias, M. (2004). Molecular physiology and modulation of somatodendritic A-type potassium channels. Molecular and cellular neurosciences 27, 343–369.

Kaeser, P.S., and Regehr, W.G. (2014). Molecular mechanisms for synchronous, asynchronous, and spontaneous neurotransmitter release. Annual review of physiology 76, 333–363.

Kaeser, P.S., and Regehr, W.G. (2017). The readily releasable pool of synaptic vesicles. Current opinion in neurobiology 43, 63–70.

Kavalali, E.T. (2015). The mechanisms and functions of spontaneous neurotransmitter release. Nature reviews Neuroscience 16, 5–16.

Korn, H., and Faber, D.S. (1991). Quantal analysis and synaptic efficacy in the CNS. Trends in neurosciences 14, 439–445.

Kougioumtzidou, E., Shimizu, T., Hamilton, N.B., Tohyama, K., Sprengel, R., Monyer, H., Attwell, D., and Richardson, W.D. (2017). Signalling through AMPA receptors on oligodendrocyte precursors promotes myelination by enhancing oligodendrocyte survival. eLife 6.

Kukley, M., Capetillo-Zarate, E., and Dietrich, D. (2007). Vesicular glutamate release from axons in white matter. Nature neuroscience 10, 311–320.

Kukley, M., Kiladze, M., Tognatta, R., Hans, M., Swandulla, D., Schramm, J., and Dietrich, D. (2008). Glial cells are born with synapses. FASEB journal : official publication of the Federation of American Societies for Experimental Biology 22, 2957–2969.

Kukley, M., Nishiyama, A., and Dietrich, D. (2010). The fate of synaptic input to NG2 glial cells: neurons specifically downregulate transmitter release onto differentiating oligodendroglial cells. The Journal of neuroscience : the official journal of the Society for Neuroscience 30, 8320–8331.

Kula, B., Chen, T.J., and Kukley, M. (2019). Glutamatergic signaling between neurons and oligodendrocyte lineage cells: Is it synaptic or non-synaptic? Glia 67, 2071–2091.

Kula, B., and Kukley, M. (2025). Geometric principles determining the morphology of oligodendrocyte precursor cells in brain white matter. BioRxiv.

Larson, V.A., Zhang, Y., and Bergles, D.E. (2016). Electrophysiological properties of NG2(+) cells: Matching physiological studies with gene expression profiles. Brain research 1638, 138–160.

Lee, S.H., Kim, K.R., Ryu, S.Y., Son, S., Hong, H.S., Mook-Jung, I., Lee, S.H., and Ho, W.K. (2012). Impaired short-term plasticity in mossy fiber synapses caused by mitochondrial dysfunction of dentate granule cells is the earliest synaptic deficit in a mouse model of Alzheimer’s disease. The Journal of neuroscience : the official journal of the Society for Neuroscience 32, 5953–5963.

Lisman, J.E. (1997). Bursts as a unit of neural information: making unreliable synapses reliable. Trends in neurosciences 20, 38–43.

Liu, S.J., and Zukin, R.S. (2007). Ca2+-permeable AMPA receptors in synaptic plasticity and neuronal death. Trends in neurosciences 30, 126–134.

Mitew, S., Hay, C.M., Peckham, H., Xiao, J., Koenning, M., and Emery, B. (2014). Mechanisms regulating the development of oligodendrocytes and central nervous system myelin. Neuroscience 276, 29–47.

Mitterdorfer, J., and Bean, B.P. (2002). Potassium currents during the action potential of hippocampal CA3 neurons. The Journal of neuroscience : the official journal of the Society for Neuroscience 22, 10106–10115.

Mizuno, H., Hirano, T., and Tagawa, Y. (2010). Pre-synaptic and post-synaptic neuronal activity supports the axon development of callosal projection neurons during different post-natal periods in the mouse cerebral cortex. The European journal of neuroscience 31, 410–424.

Mozhayeva, M.G., Sara, Y., Liu, X., and Kavalali, E.T. (2002). Development of vesicle pools during maturation of hippocampal synapses. The Journal of neuroscience : the official journal of the Society for Neuroscience 22, 654–665.

Nagy, B., Hovhannisyan, A., Barzan, R., Chen, T.J., and Kukley, M. (2017). Different patterns of neuronal activity trigger distinct responses of oligodendrocyte precursor cells in the corpus callosum. PLoS biology 15, e2001993.

Oswald, A.M., and Reyes, A.D. (2008). Maturation of intrinsic and synaptic properties of layer 2/3 pyramidal neurons in mouse auditory cortex. Journal of neurophysiology 99, 2998–3008.

Pernia-Andrade, A.J., Goswami, S.P., Stickler, Y., Frobe, U., Schlogl, A., and Jonas, P. (2012). A deconvolution-based method with high sensitivity and temporal resolution for detection of spontaneous synaptic currents in vitro and in vivo. Biophysical journal 103, 1429–1439.

Pivonkova, H., Sitnikov, S., Kamen, Y., Vanhaesebrouck, A., Matthey, M., Spitzer, S.O., Ng, Y.T., Tao, C., de Faria, O., Jr., Varga, B.V., et al. (2024). Heterogeneity in oligodendrocyte precursor cell proliferation is dynamic and driven by passive bioelectrical properties. Cell reports 43, 114873.

Ramos, R.L., Tam, D.M., and Brumberg, J.C. (2008). Physiology and morphology of callosal projection neurons in mouse. Neuroscience 153, 654–663.

Redman, S. (1990). Quantal analysis of synaptic potentials in neurons of the central nervous system. Physiological reviews 70, 165–198.

Regehr, W.G. (2012). Short-term presynaptic plasticity. Cold Spring Harbor perspectives in biology 4, a005702.

Reyes, A., and Sakmann, B. (1999). Developmental switch in the short-term modification of unitary EPSPs evoked in layer 2/3 and layer 5 pyramidal neurons of rat neocortex. The Journal of neuroscience : the official journal of the Society for Neuroscience 19, 3827–3835.

Rozov, A., Zakharova, Y., Vazetdinova, A., and Valiullina-Rakhmatullina, F. (2018). The Role of Polyamine-Dependent Facilitation of Calcium Permeable AMPARs in Short-Term Synaptic Enhancement. Frontiers in cellular neuroscience 12, 345.

Sabatini, B.L., and Regehr, W.G. (1999). Timing of synaptic transmission. Annual review of physiology 61, 521–542.

Schneggenburger, R., Sakaba, T., and Neher, E. (2002). Vesicle pools and short-term synaptic depression: lessons from a large synapse. Trends in neurosciences 25, 206–212.

Slater, B.J., and Isaacson, J.S. (2020). Interhemispheric Callosal Projections Sharpen Frequency Tuning and Enforce Response Fidelity in Primary Auditory Cortex. eNeuro 7.

Spitzer, S.O., Sitnikov, S., Kamen, Y., Evans, K.A., Kronenberg-Versteeg, D., Dietmann, S., de Faria, O., Jr., Agathou, S., and Karadottir, R.T. (2019). Oligodendrocyte Progenitor Cells Become Regionally Diverse and Heterogeneous with Age. Neuron 101, 459–471 e455.

Spruston, N., Jaffe, D.B., and Johnston, D. (1994). Dendritic attenuation of synaptic potentials and currents: the role of passive membrane properties. Trends in neurosciences 17, 161–166.

Storm, J.F. (1988). Temporal integration by a slowly inactivating K+ current in hippocampal neurons. Nature 336, 379–381.

Stubblefield, E.A., and Benke, T.A. (2010). Distinct AMPA-type glutamatergic synapses in developing rat CA1 hippocampus. Journal of neurophysiology 104, 1899–1912.

Sudhof, T.C. (2012). The presynaptic active zone. Neuron 75, 11–25.

Sun, W., and Dietrich, D. (2013). Synaptic integration by NG2 cells. Frontiers in cellular neuroscience 7, 255.

Suo, N., Guo, Y.E., He, B., Gu, H., and Xie, X. (2019). Inhibition of MAPK/ERK pathway promotes oligodendrocytes generation and recovery of demyelinating diseases. Glia 67, 1320–1332.

Swanson, G.T., Kamboj, S.K., and Cull-Candy, S.G. (1997). Single-channel properties of recombinant AMPA receptors depend on RNA editing, splice variation, and subunit composition. The Journal of neuroscience : the official journal of the Society for Neuroscience 17, 58–69.

Tagawa, Y., and Hirano, T. (2012). Activity-dependent callosal axon projections in neonatal mouse cerebral cortex. Neural plasticity 2012, 797295.

Tran, V., Miki, T., and Marty, A. (2022). Three small vesicular pools in sequence govern synaptic response dynamics during action potential trains. Proceedings of the National Academy of Sciences of the United States of America 119.

Traynelis, S.F., Wollmuth, L.P., McBain, C.J., Menniti, F.S., Vance, K.M., Ogden, K.K., Hansen, K.B., Yuan, H., Myers, S.J., and Dingledine, R. (2010). Glutamate receptor ion channels: structure, regulation, and function. Pharmacological reviews 62, 405–496.

Trommershauser, J., Schneggenburger, R., Zippelius, A., and Neher, E. (2003). Heterogeneous presynaptic release probabilities: functional relevance for short-term plasticity. Biophysical journal 84, 1563–1579.

Trussell, L.O., Zhang, S., and Raman, I.M. (1993). Desensitization of AMPA receptors upon multiquantal neurotransmitter release. Neuron 10, 1185–1196.

Voronin, L.L., and Cherubini, E. (2004). ’Deaf, mute and whispering’ silent synapses: their role in synaptic plasticity. The Journal of physiology 557, 3–12.

Waggener, C.T., Dupree, J.L., Elgersma, Y., and Fuss, B. (2013). CaMKIIbeta regulates oligodendrocyte maturation and CNS myelination. The Journal of neuroscience : the official journal of the Society for Neuroscience 33, 10453–10458.

Wen, H., Hubbard, J.M., Rakela, B., Linhoff, M.W., Mandel, G., and Brehm, P. (2013). Synchronous and asynchronous modes of synaptic transmission utilize different calcium sources. eLife 2, e01206.

Williams, S.R., and Mitchell, S.J. (2008). Direct measurement of somatic voltage clamp errors in central neurons. Nature neuroscience 11, 790–798.

Zucker, R.S., and Regehr, W.G. (2002). Short-term synaptic plasticity. Annual review of physiology 64, 355–405.

